# Seeing isn’t believing? Mixed effects of a perspective-getting intervention to improve mentoring relationships for science doctoral students

**DOI:** 10.1101/2024.10.25.620232

**Authors:** Trevor T. Tuma, Heather N. Fedesco, Emily Q. Rosenzweig, Xiao-Yin Chen, Erin L. Dolan

**Author notes:** **Author for Correspondence**, Erin L. Dolan, Department of Biochemistry & Molecular Biology, University of Georgia, B210B Davison Life Sciences Building, Athens, GA, 30602. Co-first authors TTT & HNF contributed equally to this manuscript, and each has the right to list themselves as first author on their CVs.

## Abstract

Science doctoral students can experience negative interactions with faculty mentors and internalize these experiences, potentially leading to self-blame and undermining their research self-efficacy. Helping students perceive these interactions adaptively may protect their research self-efficacy and maintain functional mentoring relationships. We conducted a pre-registered, longitudinal field experiment of a novel perspective-getting intervention combined with attribution retraining to help students avoid self-blame and preserve research self-efficacy. Science doctoral students (*n* = 155) were randomly assigned to read about mentor perspectives on negative interactions (i.e., Perspective-getting Condition) or about mentoring with no mentor perspective (i.e., control condition). Contrary to our hypotheses, we found no main effects of the intervention on students’ self-blame or research self-efficacy. However, for students with lower pre-intervention mentorship relationship satisfaction, the intervention preserved research self-efficacy six months later. This study provides evidence that perspective getting may be protective for students who are most in need of relationship intervention.

**Educational Relevance and Implications Statement:** Effective mentoring relationships are fundamental for promoting the success of doctoral students in science, yet not all mentoring relationships are high quality. This study assessed the effectiveness of a brief perspective-getting intervention (where students are given the perspective of what it is like to be a research mentor) that aimed to protect science doctoral students from blaming themselves for negative interactions with faculty mentors and maintain their research self-efficacy. Results showed that on average across all students, the intervention did not affect students’ self-blame for negative interactions or their research self-efficacy. However, the intervention did help students with less satisfying mentoring relationships maintain their self-efficacy. Thus, perspective-getting shows some promise for protecting science doctoral students from harm that can be caused by negative interactions with faculty mentors.

## 1. INTRODUCTION

Research mentoring relationships in STEM fields are championed for their role in preparing the next generation of scientists (National Academies of Sciences, Engineering, & Medicine, 2019). The quality and success of graduate students’ educational experiences are heavily influenced by their mentoring relationships with their research advisors (Barnes & Austin, 2009; Sverdlik et al., 2018; Zhao et al., 2007). Yet, a growing body of research indicates that not all student-faculty mentoring relationships are fulfilling (Burt et al., 2019; Clark et al., 2000; Goodyear et al., 1992; Griffin et al., 2022; Limeri et al., 2019; Tuma et al., 2021). In a study by Tuma and colleagues (2021), life science doctoral students reported experiencing dysfunctional and negative mentoring experiences with their research advisors. Negative experiences with mentors included a lack of mentor accessibility, limited career or psychosocial support, and limited mentor expertise, as well as mentors engaging in abusive behaviors such as yelling, coercion, threats, deceit, and belittling (Tuma et al., 2021).

Negative mentoring experiences are uniquely and negatively predictive of undergraduates’ scientific self-efficacy, science identity, research value beliefs (e.g., intrinsic and communal value, opportunity cost), and career intentions (Limeri et al., 2024). Within the context of graduate education, negative experiences with mentors are a hypothesized contributor to the high attrition rates from STEM graduate programs and deteriorating well-being reported by graduate students (Evans et al., 2018; Sowell et al., 2015; Tuma et al., 2021). Although robust research examining the consequences of negative mentoring experiences are limited, national reports and policy recommendations have indicated that there is an urgent and compelling need to improve the quality of mentorship that graduate students’ experience (National Academies of Sciences, Engineering, & Medicine, 2018, 2019; National Science Foundation, 2024). Together, this evidence suggests that approaches aimed at improving mentoring relationship quality and supporting graduate students’ in minimizing potential consequences of these experiences are needed. In the present study, we designed and tested a novel intervention to address this goal.

Guided by research suggesting that students may blame themselves for negative interactions with their mentors (Fedesco et al., 2023), the goal of the present study was to test the efficacy of a perspective-getting intervention that leveraged attribution retraining techniques to help graduate students avoid self-blame and preserve their research self-efficacy when encountering negative interactions with their mentors.

### 1.1 Existing interventions to improve mentoring relationships

Most mentorship-focused interventions designed to date lack strong experimental evidence to evaluate the effectiveness of the intervention (Gangrade et al., 2024) or have been designed primarily for mentors (Gangrade et al., 2024; National Academies of Sciences, Engineering, & Medicine, 2019). While these mentor-specific interventions have yielded promising outcomes, (Byars-Winston et al., 2023; Lewis et al., 2017; Pfund et al., 2014, 2015), they put mentors in control of any changes. Such approaches provide limited opportunities for mentees to develop adaptive beliefs and foster their agency when facing challenging interactions with their mentor. Mentee-specific interventions can shift the focus away from relying on the mentor’s goodwill to improve the relationship and instead empower mentees to take initiative and exercise their agency in their mentoring relationship. For example, “mentoring up” aims to equip mentees with the skills to shape their mentoring relationships (Lee et al., 2015) and may improve mentorship quality (Risner et al., 2020). Yet, mentoring up programming has focused on skill-building rather than helping students develop new or different beliefs about their mentoring relationships. Moreover, mentee-targeted programming has not explicitly addressed how to think about negative interactions with mentors or reduce the potential impact of such interactions. The present study aims to address such gaps by developing and testing the efficacy of a mentee-focused intervention to minimize the consequences of negative interactions.

### 1.2 Mentoring relationships can promote or hinder self-efficacy

One of the most influential self-evaluations is one’s beliefs in their own ability to engage, exert effort, and persevere in specific tasks (Bandura, 1986). Self-efficacy, or an individual’s context-dependent belief in their ability to successfully complete a task, is a powerful predictor of their goal-setting, effort, persistence, performance, and well-being (Bandura, 1977; Choi et al., 2012; Pajares, 1996; Robbins et al., 2004; Sitzmann & Yeo, 2013). Theory (Bandura, 1997) and evidence (Byars-Winston et al., 2017; Chen & Usher, 2013; Sheu et al., 2018; Usher & Pajares, 2008) point to four main sources that form an individual’s self-efficacy beliefs: mastery experiences (i.e., their past performance), vicarious experiences (i.e., observing others’ performance), verbal and social persuasion (i.e., evaluative messages and encouragement from influential others), and affective arousal (i.e., positive and negative emotional states). Mentoring relationships offer multiple avenues to shape mentee self-efficacy via these sources (Ragins, 2011; Ragins & Verbos, 2017). For example, mentors give students activities at which students might succeed and fail, they provide examples of how students might learn to be successful in their field, and they provide students with verbal persuasion about their ability to do specific tasks throughout graduate school. Thus, it is not surprising that prior research shows a connection between students’ mentoring relationships and their self-efficacy (Aikens et al., 2016; Chemers et al., 2011; Estrada et al., 2018; Joshi et al., 2019; Limeri et al., 2024).

### 1.3 The potential protective effects of gaining perspectives in relationships

A growing number of studies have indicated that brief, theory-informed, social-psychological interventions can lead to long-lasting positive changes in individuals’ beliefs (Yeager & Walton, 2011). Psychologically precise interventions that address specific beliefs and engage individuals in recursive thought-behavior cycles (sometimes called “wise” interventions in the social-psychological literature) can promote change by targeting individuals’ need to understand, need for self-integrity, and need to belong can help improve individuals’ attitudes, behaviors, and well-being (Walton & Wilson, 2018). One such intervention that may function in this way is referred to as perspective taking. Perspective taking is an active cognitive process that involves thinking about situations from another’s perspective in order to understand their thoughts, motivations, and intentions (Ku et al., 2015). The process of perspective taking helps individuals conceptualize and relate to why others may be thinking, feeling, or behaving the way they are (Galinsky et al., 2005). Perspective taking can yield relational benefits such as liking (Blatt et al., 2010; Davis et al., 1996), closeness (Galinsky et al., 2008; Peterson et al., 2015), and empathy (Bruneau & Saxe, 2012; Okonofua et al., 2016; Zhou et al., 2017). Perspective taking can also foster prosocial behaviors, such as the provision of empathic support (Fasbender et al., 2020) and helping behavior (Grant & Berry, 2011; Shih et al., 2009). Finally, perspective taking can improve individual and group outcomes that rely on positive orientations toward others, such as team performance (Hoever et al., 2012) and job satisfaction (Parmar et al., 2023).

Perspective-taking interventions illustrate that it is possible to adjust individuals’ beliefs about their relationships using psychologically “wise” techniques, pointing to one potential pathway to help improve the quality of mentor-mentee relationships. Although research on perspective taking is encouraging, asking individuals to imagine others’ viewpoints may not result in an accurate understanding (Eyal et al., 2018). For instance, the perspective taker may be influenced by egocentric bias (i.e., their own perspective on the situation) such that they become entrenched in their own perspective, leading to greater misunderstanding and relational harm (Epley et al., 2006; Vorauer & Quesnel, 2013). Some have argued that, to understand someone’s perspective, individuals need to *get* their perspective rather than *take* it (Eyal et al., 2018). Engaging in a cognitive process where individuals are *given* the perspective of another (i.e., perspective *getting*), rather than asking them to put themselves in the other’s shoes (i.e., perspective *taking*), may lead to greater benefits (Eyal et al., 2018). Providing individuals with the actual perspectives of others may also support them in being able to comprehend, contemplate, and consider this perspective recursively in the future.

Perspective getting may be particularly promising for improving the quality of graduate student-faculty mentoring relationships. For instance, providing student mentees with faculty mentors’ perspectives may aid in their ability to understand the reasons behind negative interactions. Doing so has the potential to mitigate mentee self-blame, maintain a positive orientation toward the relationship, and assist them in remaining open to the provision of mentorship support. It is also plausible that perspective getting may reduce potential consequences of negative mentoring. For instance, negative interactions with research mentors may function as a form of negative social persuasion (Bandura, 1997; Usher & Pajares, 2008), undermining students’ research self-efficacy. Perspective getting may preserve students’ research self-efficacy by empowering them to approach challenges in their mentoring relationship adaptively, which has potential downstream implications for their well-being, scholarly productivity, and success in graduate school (Paglis et al., 2006; Zhang et al., 2022). Although perspective-getting interventions have been shown to facilitate positive relationships across a variety of contexts (Ku et al., 2015), little research has examined if perspective getting can be leveraged to enhance the quality and outcomes of mentoring relationships.

### 1.4 Using attribution retraining to enhance intervention effectiveness

To maximize the effectiveness of a perspective-getting intervention, we leveraged techniques from existing motivational interventions that help students re-attribute the causes of negative experiences in more adaptive ways (i.e., “attribution retraining” interventions; Graham & Taylor, 2022). The attribution theory of motivation (Weiner, 1985) posits that individuals’ interpretations of the cause of an event (i.e., their attributions) have strong influences on individuals’ subsequent confidence and likelihood of engaging with similar tasks in the future. Attributions are posited to have three major dimensions: the extent to which the event was caused internally or externally to the individual (the locus of causality), the extent to which the event was something that was controllable by either the individual or other sources (controllability), and the extent to which the event is likely to occur again or is a rarity (stability). Extensive research suggests that if individuals attribute challenges to causes that are internal, uncontrollable, and stable (i.e., one’s own low ability or issues with one’s personality), this undermines their self-efficacy regarding future tasks, their affect during related tasks, and their likelihood of engaging with similar tasks moving forward (Graham, 2020; Perry et al., 2010).

When applied to student-faculty mentoring relationships, attribution theory suggests that, when students blame negative interactions with their mentors on their own traits or ability, this is particularly problematic for supporting their self-efficacy and graduate school satisfaction. Given that students may blame themselves for negative interactions with their mentors (Fedesco et al., 2023), it is particularly important to address self-blame when using perspective getting techniques in order to foster effective experiences for graduate students.

Attribution-retraining (AR) interventions are thought to function by shifting perceptions of the causes of negative events to avoid self-blame (Graham & Taylor, 2022; Siegel Robertson, 2000). These interventions can change students’ attributions and promote students’ classroom well-being, grades, and effort (Hamm et al., 2020; Lazowski & Hulleman, 2016; Parker et al., 2016; Perry et al., 2010). The impacts of AR interventions have been replicated across various contexts and have led to sustained improvements detectable up to eight years following the intervention (Boese et al., 2013; Hamm et al., 2017, 2020; Haynes et al., 2006). More recently, studies have shown that AR interventions may lead to the greatest benefits for students experiencing academic struggles (Hamm et al., 2020; Perry & Hamm, 2017). AR interventions that shift how students interpret challenges may be particularly beneficial for those who are struggling because they might be particularly likely to experience challenges that they blame themselves for (Hamm et al., 2020). These adjustments to core attribution processes may lead to changes in cognition, emotion, and achievement motivation, but take time to manifest as individuals shift their beliefs about their successes and failures (Hamm et al., 2017; Parker et al., 2016).

To date, AR interventions have not been applied to the context of mentoring relationships. Such AR interventions may be beneficial for reducing the potential consequences of negative mentoring experiences because they may reduce the likelihood of self-blame and prevent declines in students’ self-efficacy. For example, a negative interaction with a research mentor may lead to lower research self-efficacy if the student blames themselves as the cause of the poor interaction. Assisting the student to perceive the interaction as being caused by a combination of factors, including ones that are external or unstable, may protect their self-efficacy amid such challenges. For this reason, AR techniques coupled with perspective getting may be particularly helpful to assist mentees with recognizing factors beyond their mentor’s control (e.g., job & personal demands) that may contribute to negative interactions. This perspective may enhance their recognition of the complexity of situations faced by their mentors, thereby reducing feelings of self-blame and ultimately maintaining their self-efficacy.

### 1.5 The present study

We conducted a randomized controlled trial (RCT) to experimentally test the efficacy of a brief perspective-getting intervention among science doctoral students. We leveraged research on mentoring, interpersonal relationships, and motivation to develop the intervention through an iterative, design-based process with science doctoral students in our target population (Figure 1). We then tested the effects of the intervention versus a control condition on doctoral students’ self-blame for negative interactions with their research mentor (i.e., their locus of causality attributions) and their research self-efficacy after the intervention. We had two preregistered hypotheses regarding how the perspective-getting intervention influenced theoretically relevant student outcomes. First, we hypothesized that doctoral students in the Perspective-getting Condition would be less likely than those in the control condition to attribute negative interactions with research mentors to internal causes immediately after receiving the intervention. Second, we hypothesized that doctoral students in the Perspective-getting Condition would report higher research self-efficacy relative to those in the control condition immediately after receiving the intervention.

**Figure 1:**
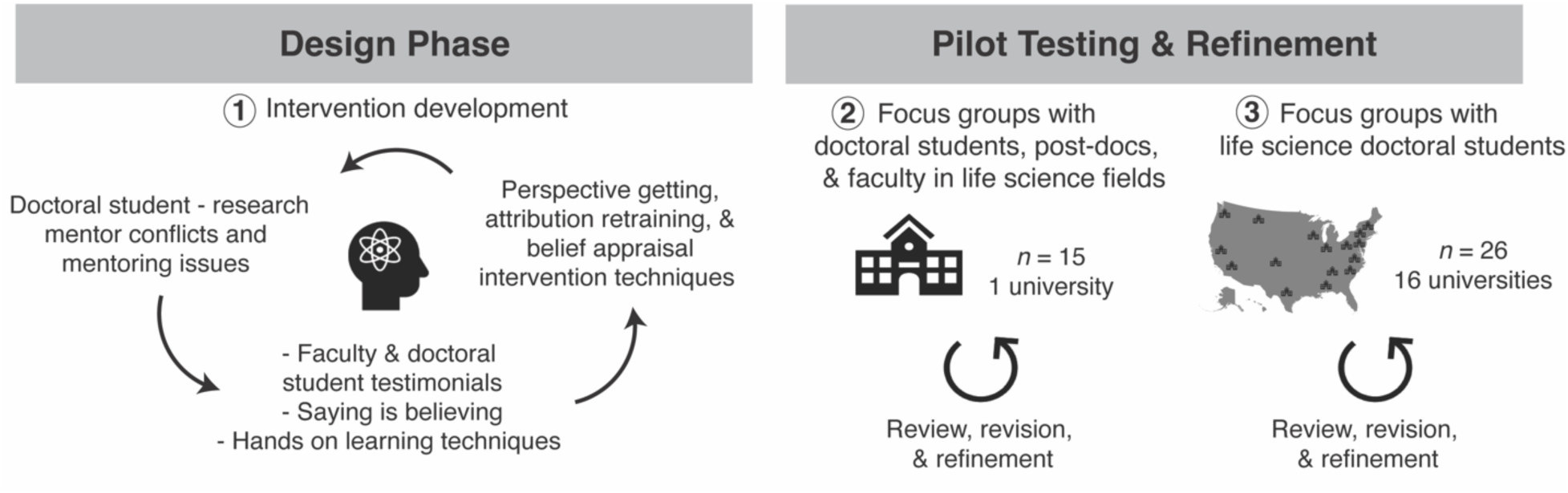
Iterative design-based process for developing the perspective-getting intervention. We developed the *Perspective-getting Condition* materials by drawing from established perspective-getting intervention approaches and research on the negative mentoring experiences of doctoral students. We designed the materials to include realistic scenarios and quotations characteristic of doctoral student-faculty mentoring relationships. We also integrated principles from attribution retraining and belief re-appraisal to influence doctoral students’ attributions about the causes of their negative mentoring experiences. We then pilot tested and iteratively revised and refined the materials with two samples to ensure their utility and authenticity prior to the randomized control trial.

Given the novel use of perspective-getting interventions in this context, we also conducted a series of exploratory analyses. Because it takes time to shift individuals’ beliefs about the causes of their successes and failures, we examined whether the perspective-getting intervention led to changes in students’ locus of causality attributions and research-self efficacy six months after the receiving the intervention. In addition, given that AR interventions may be more beneficial for individuals with lower initial achievement, we also examined if students’ satisfaction with their mentoring relationship moderated the effectiveness of the perspective-getting intervention.

## 2. METHODS

This study was reviewed and determined to be exempt by the University of Georgia Institutional Review Board (PROJECT00005263). The hypotheses, sample sizes, methods and data analysis plans for this study were preregistered (https://osf.io/dfme7/?view_only=92678c7368a5446093bb263eeffe632d) prior to data collection and analysis.

### 2.1 Participants and context

Study participants were doctoral students in their 1^st^ through 4^th^ year in the biological or chemical sciences from 24 departments at a large, research-intensive, primarily White public university in the Southeastern United States. Doctoral students were contacted by email and provided information about the study. Potential participants were instructed that the purpose of the study was to collect data on graduate students’ experiences in their doctoral programs and on materials developed to help graduate students explore potential career pathways. Participants were offered a $40 gift card for fully completing all study activities.

Doctoral students received an email invitation in Spring 2022 to participate in the study and complete a baseline survey of the constructs of interest (details below). Of 225 survey responses, 155 represented unique, consenting students who met the inclusion criteria (i.e., 1^st^ through 4^th^ year doctoral student in the biological or chemical sciences). Of the 70 responses excluded from the analyses: four did not consent to participate in the study, one was not in their 1^st^ to 4^th^ year of graduate school, 50 started the survey but did not complete any of the intervention or control activities, one did both conditions so was excluded from analysis, and 14 were duplicate responses. Thus, 155 students represent the final analytical sample (control condition = 76, intervention condition = 79); 149 students (96% of the original sample) completed Session 1, 134 students (86% of the original sample) completed Session 2, and 96 students (62% of the original sample) completed the Long-term evaluation. Self-reported participant demographic information is included in Table 1.

**Table 1:** Demographic characteristics of participants (*n* = 155)

| **Description** | ***n* (%)** |
| --- | --- |
| **Condition** |  |
| Control | 76 (49%) |
| Intervention | 79 (51%) |
| **Gender** |  |
| Man | 51 (33%) |
| Woman | 101 (65%) |
| Non-binary or prefer to self-describe | 2 (<1%) |
| Prefer not to respond | 1 (<1%) |
| **Race** |  |
| American Indian or Alaskan Native | 3 (2%) |
| Asian | 47 (31%) |
| Black or African American | 12 (8%) |
| White | 87 (58%) |
| Prefer to self-describe | 4 (2%) |
| **Ethnicity** |  |
| Hispanic or Latinx/Latine | 16 (10%) |
| Not Hispanic | 135 (87%) |
| Prefer not to respond | 4 (3%) |
| **Generation status** |  |
| First-generation college student | 39 (25%) |
| Continuing-generation college student | 116 (75%) |
| **Disability status** |  |
| Yes | 67 (43%) |
| No | 88 (57%) |
| **Year in program** |  |
| 1<sup>st</sup> | 31 (20%) |
| 2<sup>nd</sup> | 41 (27%) |
| 3<sup>rd</sup> | 31 (20%) |
| 4<sup>th</sup> | 52 (33%) |

### 2.2 Procedure

We conducted a single-blind RCT where participants were randomly assigned in a two-cell, between-participant study design involving a *Perspective-getting Condition* and a *Control Condition* (a complete description of the condition content is described further below). To balance participants’ sociodemographic characteristics between the two conditions, we created six blocking groups based on: (1) gender (i.e., man versus woman and non-binary/third gender), (2) representation status (i.e., belonging to a racial or ethnic minority group or being first in the family to go to college versus belonging to a racial or ethnic majority group and being continuing generation), and (3) program year (1^st^ and 2^nd^ year students versus 3^rd^ and 4^th^ year students). Within each group, participants were randomly sorted and alternately assigned to the *Perspective-getting Condition* (50%) or the *Control Condition* (50%). We opted for randomization at the student level instead of at the research lab or departmental level to better control for potential mentor level (i.e., mentoring competence) and organization level (i.e., departmental support for high-quality mentorship) confounding variables.

The study procedure is summarized in Figure 2. Upon study enrollment, participants completed a Baseline survey of their research self-efficacy, locus of causality attributions, mentoring relationship satisfaction, and scholarly productivity. Participants were then assigned to condition and received two doses of the condition-specific activities. One week after completing the Baseline survey, participants completed the *Perspective-getting Condition* or *Control Condition* activities (Session 1; see next section for details). One month later, they received a second dose of the *Perspective-getting Condition* or *Control Condition* activities (Session 2; see next section). After completing Session 2, participants reported their research self-efficacy and locus of causality attribution (i.e., Postintervention). Six months after completing the Postintervention, participants completed a survey (i.e., the Long-term evaluation) assessing their research self-efficacy, locus of causality attributions, mentoring relationship satisfaction, and scholarly productivity.

**Figure 2:**
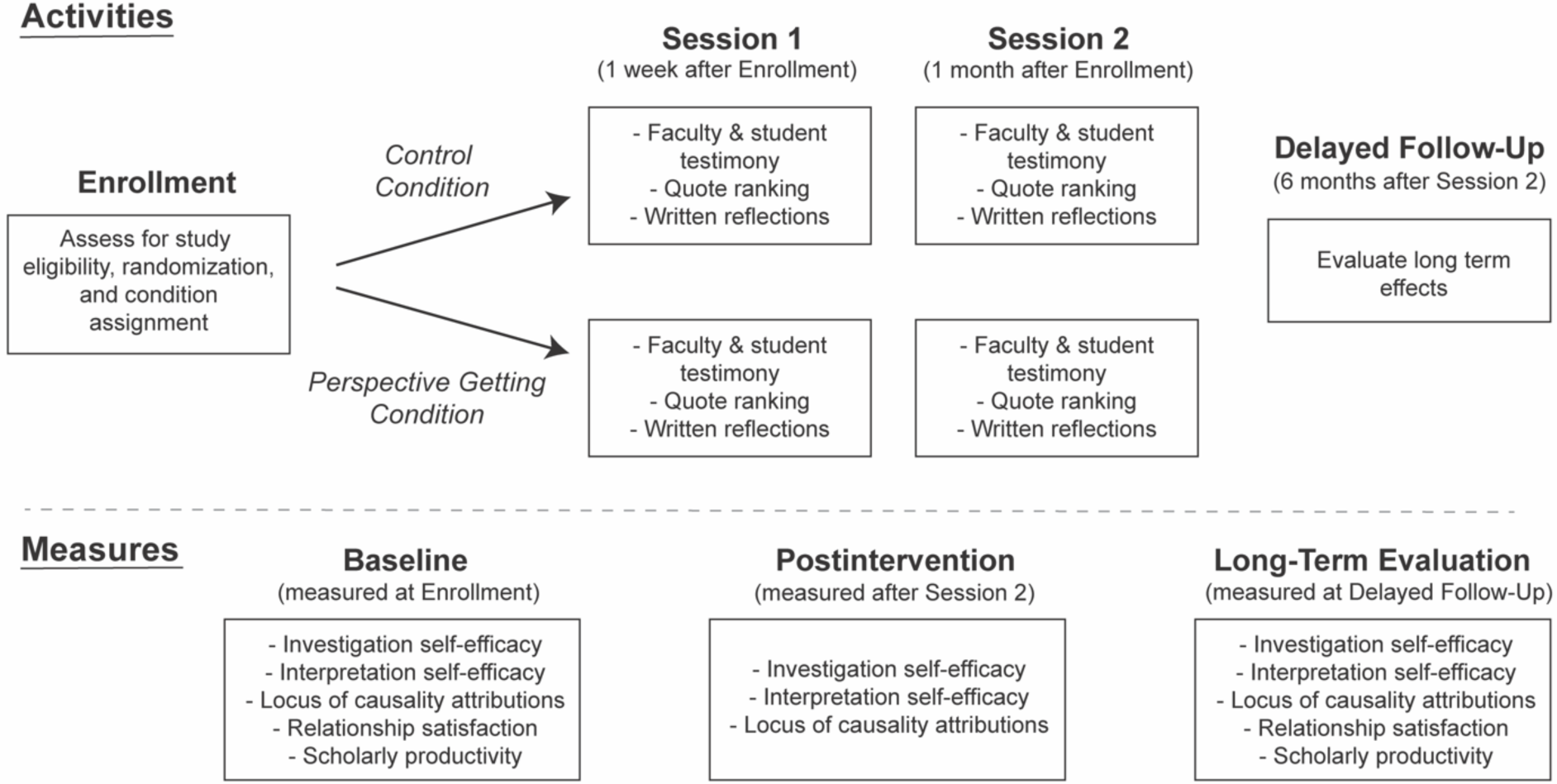
Randomized controlled trial study design process for evaluating the efficacy of the perspective-getting intervention. Biology and chemistry doctoral students enrolled in the study and completed the Baseline survey. Then, students were randomized to either the *Perspective-getting Condition*, in which they were given the perspective of what it is like to be a research mentor, or the *Control Condition*, in which they were given tips for effective mentoring. In both conditions, participants read student quotations about how other doctoral students reflected on the materials and then wrote brief written reflections on how the materials would influence them moving forward in their programs and careers during 2 independent sessions. We tested the effects of the *Perspective-getting Condition* compared to the *Control Condition* on students’ investigation and interpretation self-efficacy, locus of causality attributions, mentoring relationship satisfaction, and scholarly productivity.

### 2.3 Intervention structure, content, and development

All study activities were delivered online via the secure survey service, Qualtrics™. We elected to use an online, self-administered activity rather an in-person module to minimize participant burden of attending a face-to-face session, and because outcomes of online versus face-to-face mentoring training have been shown to be equivalent (Rogers et al., 2022). Online delivery also helped ensure a consistent and standardized experience for all participants. In both the *Perspective-getting Condition* and *Control Condition*, participants were asked to read, write, and evaluate quotations from other students. Participants in the *Perspective-getting Condition* also read and evaluated quotations from faculty members about their perspectives on interactions that had the potential to be interpreted negatively by graduate students.

We created the materials in the *Perspective-getting Condition* using a design-based process with the aim of shifting students’ attributions of the causes of negative experiences with their mentors from internal to external. In drafting the materials, we used research on negative mentoring experiences that doctoral students experience to create realistic scenarios and student quotations (Tuma et al., 2021). We also integrated faculty explanations of how their job demands influenced their mentoring practices to generate realistic faculty perspectives (Fedesco et al., 2023). Finally, we incorporated principles from attribution retraining and belief re-appraisal interventions to design the materials to change doctoral students’ attributions (Hamm et al., 2020; Rosenzweig et al., 2020, 2022; Walton & Cohen, 2007, 2011). More broadly, we designed the materials to align with recommendations about how to promote deep and active learning (Fiorella & Mayer, 2016) and optimal engagement with psychological interventions (Walton & Wilson, 2018; Yeager & Walton, 2011). The materials included opportunities for students to engage autonomously (e.g., make choices about what to discuss) and think actively and deeply about the content (e.g., complete self-reflective writing techniques, engage in self-quizzing). By leveraging this array of sources, we hoped that the initial drafts of intervention materials would (a) realistically present mentors’ perspectives on negative interactions, (b) represent concerns that doctoral students have about such interactions in a way that was relevant, and (c) prompt students to re-appraise their conflicts with mentors in a way that avoided self-blame by providing insights into situational factors that could lead to or exacerbate unpleasant interactions between students and mentors.

### 2.4 Intervention pilot testing and further refinement

After drafting the *Perspective-getting Condition* and *Control Condition* materials, we conducted pilot work to refine the study materials (Figure 1). We sought feedback on the *Perspective-getting Condition* materials through focus groups with mentees and mentors in life science fields (*n* = 15 across three groups) and life science doctoral students (*n* = 26 across six groups). Participants provided detailed feedback on content, tone, relevance, and suitability of the intervention materials. We also sought feedback on the *Control Condition* materials from graduate students and postdoctoral researchers (*n* = 6) with knowledge of STEM mentoring norms, science education, and mentorship scholarship. As a final step, two of the study authors reviewed all materials to ensure that they reflected the psychological concepts of interest and the types of mentoring experiences that we hoped to address. A complete description of iterative pilot work approaches is described in the Supplemental Materials.

### 2.5 Final intervention materials

The final versions of the *Perspective-getting Condition* and *Control Condition* materials are included in the Supplemental Materials. In Session 1 of the *Perspective-getting Condition*, participants read quotations about what it is like to be a faculty research mentor. Consistent with the attributional retraining goal of the intervention, all quotations focused on mentors’ explanations of how negative interactions with students were due to time constraints or other circumstances, not due to students themselves. To ensure that participants read and engaged with the quotations, they were asked to rate how informative each quote was and how relevant it was to their experiences as a graduate student. They were also asked to rank the quotations from most to least favorite and then to write a brief explanation about why their top ranked quote was their favorite. After reading quotations from faculty mentors, participants read three quotations from graduate students about how reading the materials from the mentors informed how the graduate attributed negative experiences with their research mentor. Participants were then asked to write 1-2 paragraphs of their own about how the information in the *Perspective-getting Condition* materials influenced how they thought about negative experiences with their own mentor both currently and as they moved forward in their graduate programs and careers. This task was again designed to encourage participant engagement and to leverage the notion of “saying-is-believing” to prompt participants to endorse the intervention message more strongly (Higgins & Rholes, 1978). In Session 2, participants were asked to write about what they remember from Session 1 (i.e., quizzing) because doing so prompts better memory of learned content (Roediger & Karpicke, 2006). Participants were then briefly reminded about the study purpose and asked to complete another session of reading faculty quotations, evaluating them, reading student examples, and writing their own quotation. These activities had the same attributional and perspective getting goals as Session 1 but used a new set of faculty and student quotations.

In the *Control Condition*, participants read strategies and tips for how to be an effective research mentor, such as how to help new mentees get oriented, how to give feedback to mentees, and how to help mentees present their research. The materials were based on established mentoring professional development materials (Pfund et al., 2014). Participants were asked to report how much they liked the information and how informative and relevant the material was for them. Participants were then asked to write reflections on how the materials might influence them as they progress in their graduate programs and careers. This activity was designed to ensure that participants engaged in thinking about mentorship but were not otherwise prompted to consider attributions or perspectives related to negative interactions with mentors. Similar to the *Perspective-getting Condition,* students in the *Control Condition* also read three quotations from graduate students who described how reading the strategies for mentoring undergraduate researchers was impactful for their experiences. As mentioned previously, none of these quotes described themes that could be construed as relating to attributions or perspective getting. Finally, participants were asked to write 1-2 paragraphs about how learning these strategies were impactful for their experiences.

### 2.6 Measures

Survey data were collected at the timepoints described in Figure 2. Brief descriptions of the variables are provided below, reliabilities are reported in Table 2, and a complete list of items is presented in the Supplemental Materials.

**Table 2:** Means, standard deviations, reliability (Ω) and zero-order correlations among study variables. *Note:* B = Baseline, P = Postintervention, L = Long-term evaluation * *p* < .05, ** *p* < .01

| Variable | *N* | *M* | *SD* | $\Omega$ | Correlations | | | | | | | | | | | |
| --- | --- | --- | --- | --- | --- | --- | --- | --- | --- | --- | --- | --- | --- | --- | --- | --- |
|  |  |  |  |  | 1 | 2 | 3 | 4 | 5 | 6 | 7 | 8 | 9 | 10 | 11 | 12 |
| 1. Investigation self-efficacy B | 155 | 4.07 | 0.63 | 0.91 | - |  |  |  |  |  |  |  |  |  |  |  |
| 2. Investigation self-efficacy P | 134 | 4.21 | 0.53 | 0.91 | .66** | - |  |  |  |  |  |  |  |  |  |  |
| 3. Investigation self-efficacy L | 104 | 4.22 | 0.51 | 0.89 | .63** | .66** | - |  |  |  |  |  |  |  |  |  |
| 4. Interpretation self-efficacy B | 155 | 3.68 | 0.69 | 0.89 | .65** | .57** | .51** | - |  |  |  |  |  |  |  |  |
| 5. Interpretation self-efficacy P | 134 | 3.87 | 0.67 | 0.89 | .50** | .76** | .60** | .69** | - |  |  |  |  |  |  |  |
| 6. Interpretation self-efficacy L | 104 | 3.91 | 0.68 | 0.90 | .52** | .57** | .74** | .55** | .64** | - |  |  |  |  |  |  |
| 7. Locus of causality attributions B | 147 | 4.31 | 1.89 | 0.70 | -.27** | -.31** | -.26* | -.24** | -.24** | -.07 | - |  |  |  |  |  |
| 8. Locus of causality attributions P | 126 | 4.52 | 1.86 | 0.79 | -.14 | -.21* | -.26** | -.15 | -.17 | -.16 | .26** | - |  |  |  |  |
| 9. Locus of causality attributions L | 93 | 4.04 | 1.91 | 0.85 | -.27** | -.22* | -.29** | -.28** | -.19 | -.09 | .25* | .34** | - |  |  |  |
| 10. Scholarly productivity B | 154 | 5.41 | 5.60 | - | .34** | .32** | .29** | .40** | .33** | .29** | -.06 | -.07 | -.18 | - |  |  |
| 11. Scholarly productivity L | 96 | 6.58 | 6.25 | - | .36** | .33** | .33** | .41** | .36** | .32** | -.06 | -.11 | -.17 | .88** | - |  |
| 12. Relationship satisfaction B | 155 | 4.05 | 1.21 | - | .01 | -.02 | .11 | -.01 | .03 | .06 | .15 | .29** | .08 | .00 | -.03 | - |
| 13. Relationship satisfaction L | 96 | 4.10 | 1.30 | - | .06 | -.00 | -.00 | .02 | .07 | .07 | .13 | .16 | .08 | .14 | .09 | .42** |

#### 2.6.1 Locus of causality attributions for negative interactions

Locus of causality attributions about negative interactions with faculty mentors were measured at the Baseline, Postintervention, and Long-term evaluation timepoints using a causal attributions scale adapted from McAuley and colleagues (1992). Consistent with our preregistered hypotheses, students were prompted to recall a negative or unpleasant experience with their mentor, write 2-3 sentences about the experience and how it made them feel. They then responded to three items on a 9-point semantic differential scale whether the cause of the experience was internal or external to themselves (locus of causality; 1 = “something that reflects an aspect of the situation”; 9 = “something that reflects an aspect of yourself”). The scale demonstrated acceptable internal reliability at all timepoints (Table 2).

#### 2.6.2 Interpretation & investigation research self-efficacy

Research self-efficacy was measured at the Baseline, Postintervention, and Long-term evaluation timepoints using a scale adapted from Kardash (2000). Our measurement model results indicated that this scale was best represented by two subscales (see Supplemental Materials): investigation self-efficacy and interpretation self-efficacy. We used six items to measure investigation self-efficacy, or students’ confidence in their abilities to design and conduct investigations (A sample item is: *Please rate your confidence in your ability to understand contemporary concepts in your field)*. We also used six items to measure students’ interpretation self-efficacy, or confidence in their abilities to make sense of and communicate scientific results (A sample item is: *Please rate your confidence in your ability to orally communicate the results of research projects*). Students were asked to rate their confidence on a five-point, Likert-type scale (i.e., 1 = “not at all confident”; 5 = “very confident”). The scale demonstrated high internal reliability for both investigative and interpretation research self-efficacy at all timepoints (Table 2).

#### 2.6.3 Mentoring relationship satisfaction

Mentoring relationship satisfaction was measured at the Baseline and Long-term evaluation timepoints using a single item (“Overall, how satisfied are you with the quality of your relationship with your graduate research mentor?”) and a five-point, Likert-type scale (i.e., 1 = “very dissatisfied”; 5 = “very satisfied”).

#### 2.6.4 Scholarly productivity

Scholarly productivity was measured at the Baseline and Long-term evaluation timepoints by asking participants to report counts of first-author publications, co-authored publications, and conference talks/posters presentations they had delivered. These counts were summed to create a score of scholarly productivity representing the number of times they created a scholarly product.

### 2.7 Psychometric analyses

Assessment of the measurement models is reported in depth in the Supplemental Materials. In brief, we conducted confirmatory factor analysis (CFA) for our measures of self-efficacy and attributions of negative interactions. Consistent with best practices, we examined the adequacy of data-model fit using a variety of fit indices, including the χ^2^ test, RMSEA, CFI, TLI, and SRMR (Hu & Bentler, 1999). The model fit indices indicated good fit for the attributions scale but misfit for the research self-efficacy scale, which then required re-specification as detailed in the Supplemental Materials. We used the re-specified scale in all subsequent analyses.

### 2.8 Intervention adherence

We verified intervention adherence (i.e., the extent to which students completed materials as intended) using several pieces of evidence. Specifically, we examined (1) completion of activities in both conditions, (2) engagement time spent completing the intervention materials, and (3) coding of students’ self-generated descriptions of a negative interaction with their research mentor. A full description of each metric are reported in the Supplemental Materials.

### 2.9 Analytic plan

We followed the analytical plan noted in the preregistration (https://osf.io/dfme7/?view_only=92678c7368a5446093bb263eeffe632d) with two minor deviations. We describe the differences between our planned and reported analyses in the Supplemental Materials.

We used linear regression to test our two preregistered hypotheses and evaluate the effectiveness of the perspective-getting intervention on self-efficacy and locus of causality attributions at the Postintervention time point. Our linear analytical model included two terms: a comparison code representing condition (intervention coded as 1, control coded as 0) and a covariate of program year (scaled from 1-4). All regression analyses were performed in MPlus version 8, with full information maximum likelihood estimation to account for missing data. A sensitivity analysis conducted with G*Power (Faul et al., 2007) suggested that a sample of *n* = 155 (two tailed, *t*-test, 0.80 power, and alpha of 0.05) was sensitive enough to detect group differences with an effect size of Cohen’s *d* = 0.45.

We tested two additional exploratory hypotheses that were not preregistered. First, we separately regressed students’ investigation self-efficacy, interpretation self-efficacy, locus of causality attributions, mentoring relationship satisfaction, and scholarly productivity at the Long-term evaluation timepoint, onto the condition while controlling for program year. Second, we tested mentoring relationship satisfaction (standardized) as a moderator of treatment effects on investigation self-efficacy, interpretation self-efficacy, and locus of causality attributions at both the Postintervention and Long-term evaluation timepoints.

## 3. RESULTS

Descriptive statistics and zero-order correlations between variables are reported in Table 2, and descriptive statistics by condition (i.e., *Perspective-getting Condition* and *Control Condition*) are reported in Table 3. All variables were in expected ranges and directions as described in the Supplemental Materials. Results revealed no significant pre-intervention differences between the *Perspective-getting Condition* and *Control Condition* for all variables (see Supplemental Materials).

**Table 3:** Descriptive statistics by study condition from Baseline.

| **Variable** | *Control Condition*<br>(*n* = 76) |  | *Perspective-getting*<br>*Condition*<br>(*n* = 79) |  |
| --- | --- | --- | --- | --- |
|  | *M* | *SD* | *M* | *SD* |
| Investigation self-efficacy | 4.02 | 0.68 | 4.11 | 0.57 |
| Interpretation self-efficacy | 3.64 | 0.67 | 3.71 | 0.71 |
| Locus of causality attributions | 4.27 | 1.85 | 4.34 | 1.94 |
| Mentor relationship satisfaction | 3.91 | 1.00 | 4.18 | 1.36 |
| Scholarly productivity | 5.55 | 6.34 | 5.25 | 4.73 |

*Note:* Results indicated no significant pre-intervention differences between conditions for the variables included in the study

### 3.1 Results of preregistered analyses

Results of the regression models predicting research self-efficacy and the locus of causality attributions while controlling for year in program at the Postintervention time point are reported in Table 4. Contrary to our preregistered hypotheses, the *Perspective-getting Condition* did not have a significant main effect on participants’ investigation self-efficacy, β = −0.06, *p* = .427, interpretation self-efficacy, β = −0.08, *p* = .360, nor their locus of causality attributions, β = 0.09, *p* = .293, at Postintervention.

**Table 4:**
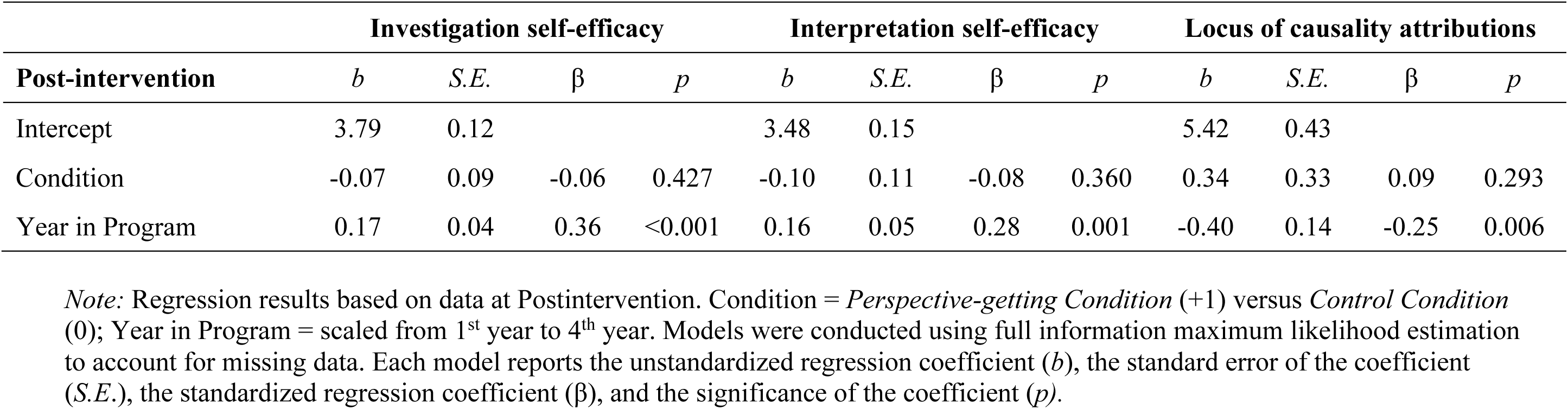
Regression results predicting investigation self-efficacy, interpretation self-efficacy, and locus of causality attributions at Postintervention.

### 3.2 Exploratory analyses of intervention effects

In exploratory analyses, we tested if the *Perspective-getting Condition* was influential in the long-term by examining students’ research-self efficacy and their locus of causality attributions six months following the intervention at the Long-term evaluation time point (Table 5). Again, the *Perspective-getting Condition* did not significantly affect participants’ investigation self-efficacy, β = −0.02, *p* = .873, interpretation self-efficacy, β = −0.08, *p* = .427, locus of causality attributions, β = −0.01, *p* = .915, mentoring relationship satisfaction, β = −0.02, *p* = .884, or scholarly productivity, β = −0.01, *p* = .908.

**Table 5:** Regression results predicting investigation self-efficacy, interpretation self-efficacy, locus of causality attributions, scholarly productivity, and relationship satisfaction at Long-term evaluation.

| **Long-term evaluation** | **Investigation self-efficacy** |  |  |  | **Interpretation self-efficacy** |  |  |  | **Locus of causality attributions** |  |  |  |
| --- | --- | --- | --- | --- | --- | --- | --- | --- | --- | --- | --- | --- |
| | *b* | *S.E.* | $\beta$ | *p* | *b* | *S.E.* | $\beta$ | *p* | *b* | *S.E.* | $\beta$ | *p* |
| Intercept | 4.04 | 0.13 |  |  | 3.51 | 0.16 |  |  | 4.23 | 0.53 |  |  |
| Condition | -0.02 | 0.10 | -0.02 | 0.873 | -0.09 | 0.12 | -0.08 | 0.427 | -0.04 | 0.40 | -0.01 | 0.915 |
| Year in Program | 0.07 | 0.04 | 0.16 | 0.108 | 0.14 | 0.05 | 0.26 | 0.008 | -0.07 | 0.18 | -0.04 | 0.707 |

| **Long-term evaluation** | **Scholarly productivity** |  |  |  | **Relationship satisfaction** |  |  |  |
| --- | --- | --- | --- | --- | --- | --- | --- | --- |
| | *b* | *S.E.* | $\beta$ | *p* | *b* | *S.E.* | $\beta$ | *p* |
| Intercept | 4.24 | 1.68 |  |  | 4.19 | 0.28 |  |  |
| Condition | -0.15 | 1.26 | -0.01 | 0.908 | -0.03 | 0.21 | -0.02 | 0.884 |
| Year in Program | 0.91 | 0.56 | 0.17 | 0.101 | -0.06 | 0.09 | -0.07 | 0.525 |

*Note:* Regression results based on data at Long-term evaluation. Condition = *Perspective-getting Condition* (+1) versus *Control Condition* (0); Year in Program = scaled from 1<sup>st</sup> year to 4<sup>th</sup> year. Models were conducted using full information maximum likelihood estimation to account for missing data. Each model reports the unstandardized regression coefficient (*b*), the standard error of the coefficient (*S.E.*), the standardized regression coefficient ( $\beta$ ), and the significance of the coefficient (*p*).

We also tested whether mentoring relationship satisfaction moderated the effect of the *Perspective-getting Condition* on students’ investigation self-efficacy, interpretation self-efficacy, and locus of causality attributions. Although we found no moderating effect of mentoring relationship satisfaction at the Postintervention timepoint (Table 6), mentoring relationship satisfaction had a significant interaction with the *Perspective-getting Condition* on both investigation self-efficacy, β = −0.32, *p* = .026, and interpretation self-efficacy, β = −0.37, *p* = .008 at the Long-term evaluation timepoint. Students in the *Perspective-getting Condition* who reported comparatively lower levels of mentoring relationship satisfaction (i.e., estimated at 1 SD below the standardized mean score for this variable) also reported higher levels of both investigation and interpretation self-efficacy at the Long-term evaluation compared to students in the *Control Condition* who reported lower initial levels of mentoring relationship satisfaction (Figure 3). However, students who reported higher relationship satisfaction prior to the intervention (i.e., estimated at 1 SD above the standardized mean score) reported lower levels of investigation and interpretation self-efficacy at the Long-term evaluation compared to those in the *Control Condition* who reported higher initial levels of mentoring relationship satisfaction (Figure 3). At the Long-term evaluation, mentoring relationship satisfaction did not moderate the relationship between the intervention contrast and students’ locus of causality attributions, β = −0.06, *p* = .720. These findings indicate that the *Perspective-getting Condition* may benefit students’ self-efficacy who have less satisfying mentoring relationships six months after the intervention and may be costly for those with more satisfying mentoring relationships. However, this effect does not appear to be due to shifts in students’ attributions.

**Figure 3:**
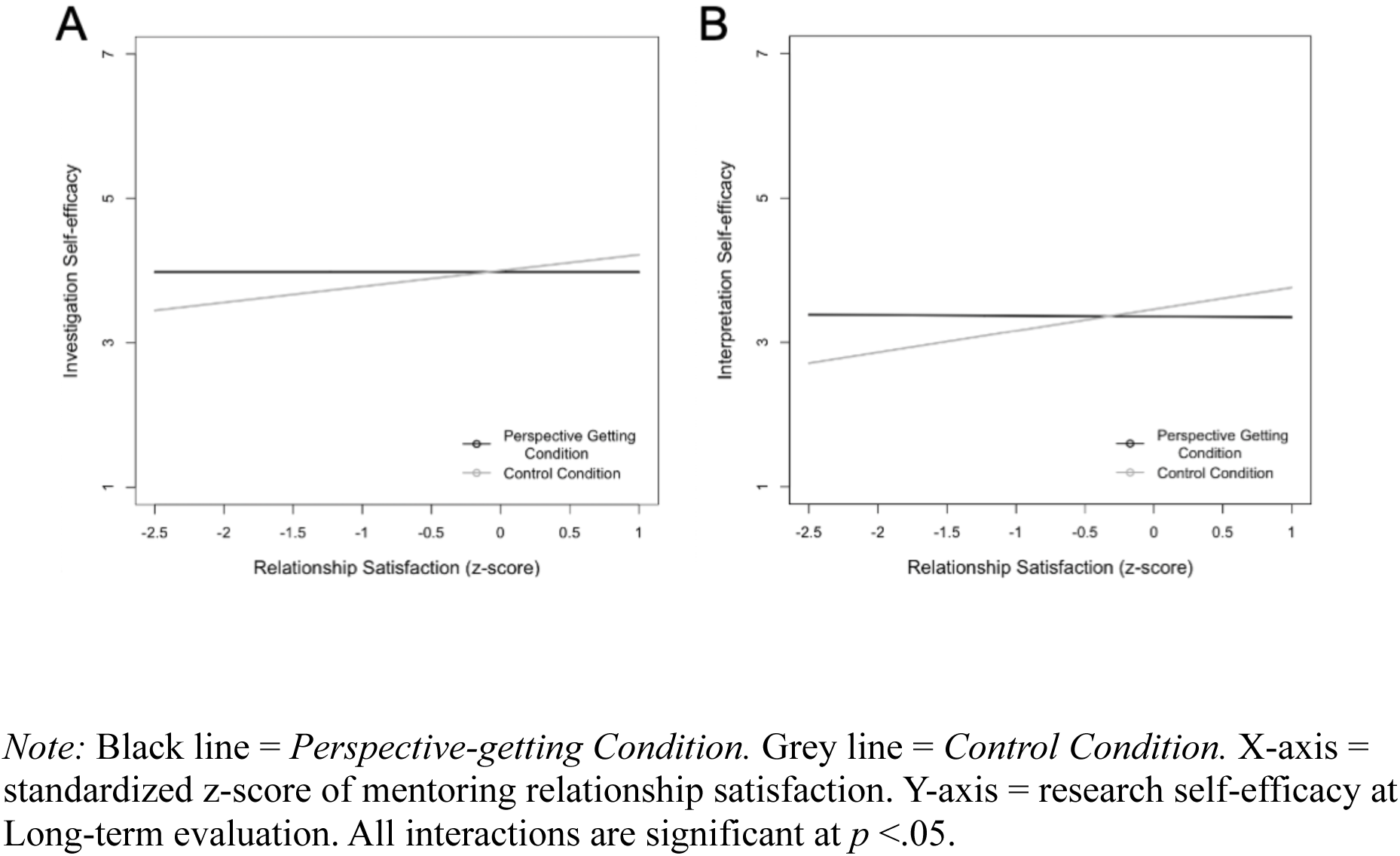
Long-term differential effects of perspective-getting dependent on students’ mentoring relationship satisfaction. *Note:* Black line = *Perspective-getting Condition.* Grey line = *Control Condition.* X-axis = standardized z-score of mentoring relationship satisfaction. Y-axis = research self-efficacy at Long-term evaluation. All interactions are significant at *p* <.05.

**Table 6:**
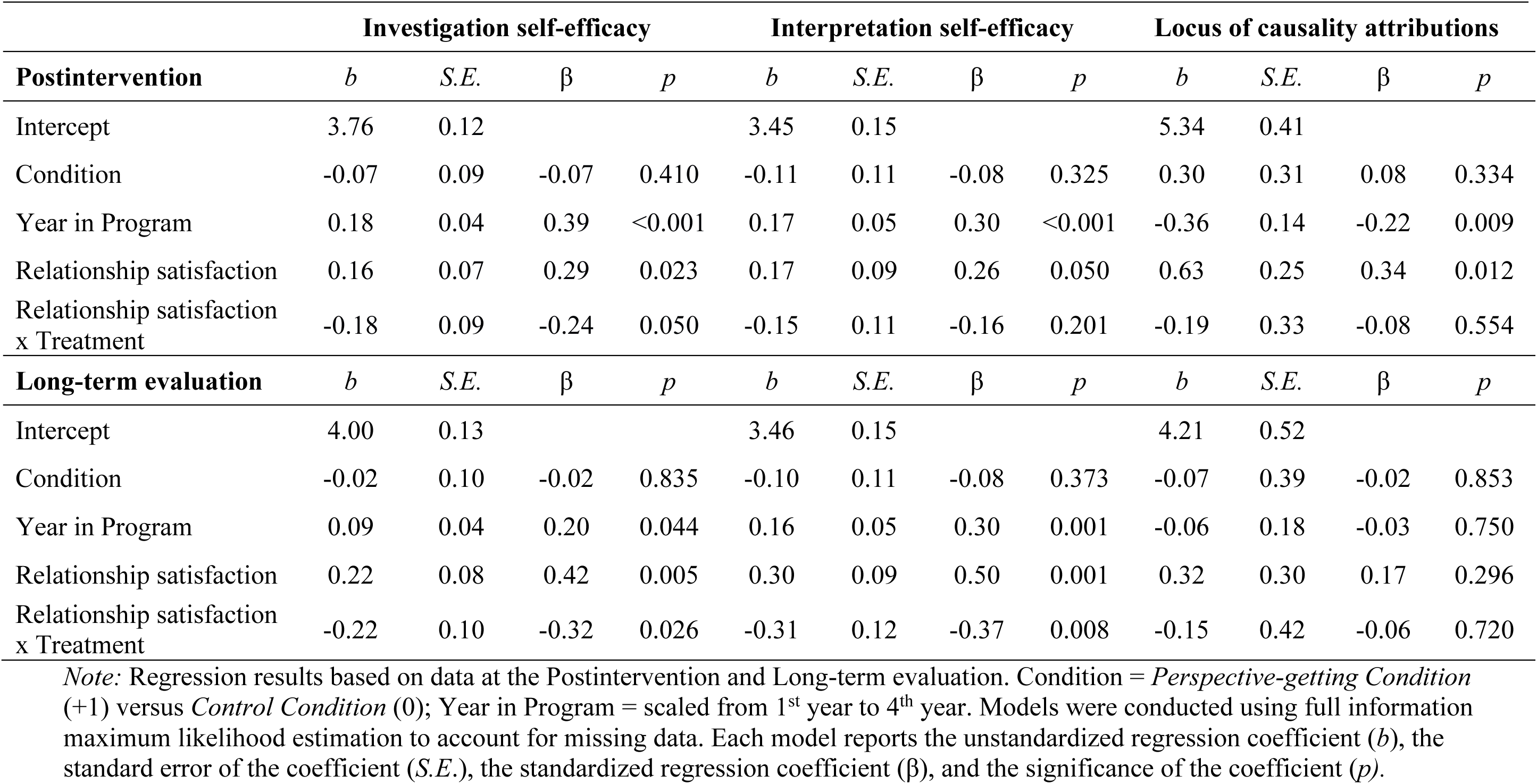
Regression results for the effect of mentoring relationship satisfaction on the relationship between the intervention and investigation self-efficacy, interpretation self-efficacy, locus of causality attributions.

|  | Investigation self-efficacy |  |  |  | Interpretation self-efficacy |  |  |  | Locus of causality attributions |  |  |  |
| --- | --- | --- | --- | --- | --- | --- | --- | --- | --- | --- | --- | --- |
| **Postintervention** | *b* | *S.E.* | $\beta$ | *p* | *b* | *S.E.* | $\beta$ | *p* | *b* | *S.E.* | $\beta$ | *p* |
| Intercept | 3.76 | 0.12 |  |  | 3.45 | 0.15 |  |  | 5.34 | 0.41 |  |  |
| Condition | -0.07 | 0.09 | -0.07 | 0.410 | -0.11 | 0.11 | -0.08 | 0.325 | 0.30 | 0.31 | 0.08 | 0.334 |
| Year in Program | 0.18 | 0.04 | 0.39 | <0.001 | 0.17 | 0.05 | 0.30 | <0.001 | -0.36 | 0.14 | -0.22 | 0.009 |
| Relationship satisfaction | 0.16 | 0.07 | 0.29 | 0.023 | 0.17 | 0.09 | 0.26 | 0.050 | 0.63 | 0.25 | 0.34 | 0.012 |
| Relationship satisfaction<br>x Treatment | -0.18 | 0.09 | -0.24 | 0.050 | -0.15 | 0.11 | -0.16 | 0.201 | -0.19 | 0.33 | -0.08 | 0.554 |
| **Long-term evaluation** | *b* | *S.E.* | $\beta$ | *p* | *b* | *S.E.* | $\beta$ | *p* | *b* | *S.E.* | $\beta$ | *p* |
| Intercept | 4.00 | 0.13 |  |  | 3.46 | 0.15 |  |  | 4.21 | 0.52 |  |  |
| Condition | -0.02 | 0.10 | -0.02 | 0.835 | -0.10 | 0.11 | -0.08 | 0.373 | -0.07 | 0.39 | -0.02 | 0.853 |
| Year in Program | 0.09 | 0.04 | 0.20 | 0.044 | 0.16 | 0.05 | 0.30 | 0.001 | -0.06 | 0.18 | -0.03 | 0.750 |
| Relationship satisfaction | 0.22 | 0.08 | 0.42 | 0.005 | 0.30 | 0.09 | 0.50 | 0.001 | 0.32 | 0.30 | 0.17 | 0.296 |
| Relationship satisfaction<br>x Treatment | -0.22 | 0.10 | -0.32 | 0.026 | -0.31 | 0.12 | -0.37 | 0.008 | -0.15 | 0.42 | -0.06 | 0.720 |

*Note:* Regression results based on data at the Postintervention and Long-term evaluation. Condition = *Perspective-getting Condition* (+1) versus *Control Condition* (0); Year in Program = scaled from 1<sup>st</sup> year to 4<sup>th</sup> year. Models were conducted using full information maximum likelihood estimation to account for missing data. Each model reports the unstandardized regression coefficient (*b*), the standard error of the coefficient (*S.E.*), the standardized regression coefficient ( $\beta$ ), and the significance of the coefficient (*p*).

### 3.3 Analysis of open-ended responses to explore potential mixed intervention effects

To understand why the *Perspective-getting Condition* had no overall effects, we examined students’ written responses about the *Perspective-getting Condition* material. Through this exploratory analysis, we sought to understand whether students internalized an understanding of mentors’ perspectives on negative interactions as we had intended. Two of the study authors qualitatively analyzed students’ text descriptions of how understanding their mentors’ perspectives influenced their perceptions of negative experiences with their mentor. To accomplish this, we reviewed student text responses and made analytic memos regarding our impressions of themes in the data. We found that student responses to the *Perspective-getting Condition* could broadly be grouped into three main categories: acceptors, rejectors, and those with mixed reactions. Students whose responses were generally positive about understanding faculty perspectives on negative interactions or stated the *Perspective-getting Condition* was useful for them were coded as ‘acceptors.’ Students whose responses suggested they did not buy in to the messages in the *Perspective-getting Condition* or wrote that the materials were not useful or applicable for their situations were coded as ‘rejectors.’ Students whose responses contained elements of both buying into and pushing back on mentor perspectives were coded as ‘mixed.’ Two researchers independently coded each participant’s response as acceptor, rejector, or mixed. Given the exploratory nature of this phase of analyses, which aimed to gain insight on how students were reacting to the *Perspective-getting Condition*, we did not systematically code open-ended responses to consensus.

We coded a total of 142 open-ended text responses from both Session 1 and Session 2 into acceptors, rejectors, or mixed responses to the *Perspective-getting Condition*, as detailed in Table 7. Six responses were non-sensical or unrelated to the prompt and were excluded from further analyses, resulting in a final analytic sample of 136 text responses. Most student responses (102, or 75%) suggested they accepted the messages in the *Perspective-getting Condition*, reporting that the materials helped them better understand their mentors’ perspectives and relate to the causes of negative interactions with their mentors. A smaller proportion of students had mixed responses (28, or 20%), which indicated that they understood mentors’ perspectives during negative interactions but were unable to fully accept or agree with them (see ‘rejectors’ below). Their responses acknowledged the challenges mentors face, while questioning whether mentors were justified in actively negatively toward students as a result.

**Table 7:** Results of coding text responses (total *n* = 136) in the *Perspective-getting Condition* materials.

| **Coded participant response** | **$n$**<br>**(%)** | **Excerpt from participant written response** |
| --- | --- | --- |
| **Acceptors:**<br>Positive response and/or reported the intervention materials as useful | 102<br>(75%) | *“Reading the mentor's tale of professional life was encouraging and reassuring. I liked their humbleness toward their students' work and also a kind of remorse for not fulfilling their duties toward the students. It made me realize that mentors try their best to be supportive of the students. Mentors also feel remorse if they cannot fulfill their duties as an mentor. Similarly, reading their excerpts was revealing as they mentioned that failing at a few things does not mean you fail in your professional life. They seem to be considerate of students more than I thought they were. Overall, it feels good to know that they do not take things personally and that what they do is mostly directed toward the betterment of the student's professionalism. These readings had a positive impact on me and on my beliefs about academia.”*<br>*“It is certainly challenging to sometimes feel like your mentor is not giving you as much attention as other students or other lab members. However, reading these quotes makes me realize it's not a conscious decision. In the past, I have been frustrated with the amount of guidance I received and sometimes have had conflict with my mentor because I felt they weren't doing exactly what I needed. However, reading these quotes makes me realize I am only a very small part of their professional life - they have a lot more on their plate (including working with other students) to work on than just appeasing all of my needs.”* |
| **Mixed:**<br>Response contained elements of both buying into and pushing back against the materials | 28<br>(20%) | *“I found it interesting to read about some people's thought process behind their mentoring decisions. I find it helpful to think about moments when I wish I had received more clear guidance as my mentor is trying to find the right balance between protecting me from mistakes and allowing me independence rather than just him not paying close enough attention. I have heard a lot about how difficult it is to be an mentor and how the demands of the job are near impossible to meet. I typically have a hard time sympathizing with these perspectives because I have seen so many examples of mentors treating their students poorly and using the stresses of the job as an excuse. I also struggle to sympathize because mentors have so much more power, receive markedly better compensation, and much more* |
|  |  | *protection than graduate students. I find it much easier to sympathize with the struggles of mentors when those struggles are related to doing their best for their students.”* |
|  |  | *“While it is helpful to understand the perspective of [mentors] better, I still believe that the selected perspectives are the minority in academia. Many [mentors] do not care about researcher development as much as they care about the next paper, the next conference, the next grant. I do not find myself particularly swayed to sympathy to [mentors] about how hard it is to mentor students.”* |
| **Rejectors:**<br>Negative<br>response and/or<br>reported the<br>intervention<br>materials as not<br>useful | 6<br>(5%) | *“Fundamentally, these responses reinforce that research mentors are human, but what's missing in those discussions is recognition that graduate students are human too. There's a power imbalance between mentors and students. Failing to acknowledge this imbalance leads to exploitation of graduate students. That highlights a key failure of academia as a whole. Students and postdocs often go unacknowledged in the success of a mentor. Yet it is often their work that serves as the foundation for getting grants, publications, and more. The responses mostly indicate why mentors fail to engage in mentorship. [But] there are many stories of students who aren't being advised yet are being exploited. There are many stories of abusive mentors who remain protected by the system.”*<br>*“I understand the challenges of being a mentor, as challenges come with all positions. However, these are not new and are part of the job. Everyone has a life outside of work to balance their time with, everyone has struggles with time management, and everyone wants to feel valued. Reading about mentor’s playing victim actually makes me feel worse about the situation. As a graduate student, we are underpaid and undervalued... To have a mentor say they are too busy to help the students that they accepted into their lab under the agreement of a training program shows they are not equipped to manage.”* |

Only a small proportion of student responses (6, or 5%) actively pushed back against the messages in the *Perspective-getting Condition* and were therefore coded as ‘rejectors’. These students reported that the material did not change their perceptions of negative interactions with mentors. For instance, participants described how they were expected to understand their mentor’s viewpoint, yet their own perspectives were not acknowledged or considered by their mentor. Students explained that this dynamic made the relationship feel one-sided and unilateral. Other ‘rejector’ responses from students indicated that the scenarios were not salient to their situations and did not reflect the severity of negative interactions they experienced in their mentoring relationships. These results indicated that most participants internalized the messages in the *Perspective-getting Condition* but approximately at least a quarter of the sample expressed at least some skepticism towards the messages. It is plausible that the intervention did not have a main effect because of varied internalization of the *Perspective-getting Condition* messages.

## 4. DISCUSSION

National calls and policy decisions have emphasized the need to improve mentorship that STEM doctoral students experience during their graduate education (National Academies of Sciences, Engineering, & Medicine, 2018, 2019; National Science Foundation, 2024). Yet, robust evaluations of approaches for improving mentoring relationships are rare (Gangrade et al., 2024). Moreover, few interventions that have been developed to protect students from potential consequences of negative interactions with faculty mentors. Here, we aimed to understand if an intervention focused on perspective getting could help students make more adaptive attributions of negative interactions with their faculty mentors and promote their research self-efficacy.

Evidence about the effectiveness of the perspective-getting intervention largely did not support our hypotheses, but our observed mixed effects point to important key boundary conditions on when this type of intervention may or may not be beneficial for doctoral students. Findings also revealed key areas where future research can continue to address this important topic. We discuss the implications and broader impacts of these findings below.

### 4.1 Extending perspective getting to mentoring relationships

Students in our study who received the *Perspective-getting Condition* had no overall differences in their causal attributions or research self-efficacy compared to students in the *Control Condition.* These results are contrary to our hypotheses and outcomes of some field studies of perspective focused interventions (e.g., Galinsky et al., 2005; Ku et al., 2015), However, these results are consistent with prior studies demonstrating that perspective focused interventions can have heterogenous effects in some cases (Tarrant et al., 2012) and negative effects in others (Epley et al., 2006; Eyal et al., 2018; Vorauer et al., 2009; Vorauer & Quesnel, 2013).

There are two possible reasons why the perspective-getting intervention did not have its desired effects overall. First, some evidence indicates that perspective focused interventions may be less efficacious in environments characterized by competition and power differentials (Todd & Galinsky, 2014). In competitive contexts, individuals may be motivated to protect themselves (Epley et al., 2006) and thus less inclined to accept others’ perspectives or take them into consideration. In addition, encouraging individuals in low-power groups to take the perspective of high-power groups may be perceived as threatening and aversive (Bruneau & Saxe, 2012).

Thus, it may be that the power differential between doctoral students and their faculty mentors, particularly in science disciplines, which are characterized by competitiveness, limited the potential effectiveness of a perspective-getting intervention. Our exploratory analysis of students’ written responses is consistent with this interpretation; a quarter of our sample expressed skepticism about or pushed back against the intervention messages. In addition, some students seemed unable to agree with the intervention’s message completely because their mentor was not asked to understand their perspective. Gaining an understanding of a mentor’s perspective may not be effective for students who feel solely responsible for improving their relationship, especially if they believe their mentor is not equally invested. Consequently, it may be that these students are less willing or motivated to understand their mentor’s viewpoint through perspective getting.

Another possibility is that many students in our study had largely satisfying relationships with their mentors and thus had little room for improvement. It may be that the students who are more satisfied with their mentoring relationships are more willing to participate in a mentoring-related study (Tuma et al., 2021). Satisfying mentoring relationships are often characterized by frequent interaction and communication (Lankau & Scandura, 2002; Liang et al., 2008), which may in turn offer opportunities for disclosure by both mentors and mentees, leading to greater mutual understanding and less need to actively get one another’s perspective without intervention. It is also likely that students who reported being more satisfied with their mentoring relationships experience fewer negative interactions with their mentor, which may result in fewer maladaptive behaviors that could be ameliorated by perspective getting.

### 4.2 Benefits for students most in need of a relationship intervention

Perhaps the most noteworthy finding of the current study is that students with lower relationship satisfaction – those who may be most in most need of a relationship intervention – reported greater self-efficacy six months after receiving the *Perspective-getting Condition* relative to those in the *Control Condition*. It is plausible that the motivational benefits of the intervention were gradual because it takes time for behavioral, cognitive, and emotional adjustments to develop. This is consistent with the principles of other “wise” psychological interventions, in that recursive thought-action cycles over time are thought to drive intervention effectiveness rather than an immediate transformative change (Walton & Wilson, 2018). Our findings are also consistent with attribution theory, which suggests that initial adjustments to core attribution processes can gradually lead to changes in cognitions, emotions, and achievement motivation, which are reciprocally reinforced over time (Hamm et al., 2017; Parker et al., 2016). For instance, Hamm and colleagues’ (2020) motivational intervention aimed at restructuring students’ causal attributions were shown to produce behavioral changes that were sustained for up to eight years following the intervention (Hamm et al., 2020).

However, we did not observe a delayed effect on students’ causal attributions of negative interactions with mentors, which indicates that the effect on students’ self-efficacy likely was a direct result of the intervention and was not directly due to how they attribute their own negative interactions with mentors. As was noted earlier, one way that self-efficacy develops is via social persuasion and vicarious learning experiences, which may be shaped by interactions with mentors. It may be that students in more tenuous mentoring relationships receive more negative social persuasion and have less access to vicarious models of success, while students in supportive relationships do not. Perspective getting may have been beneficial by allowing students to receive social persuasion and vicarious examples of success via other faculty mentors and student quotes, helping them feel more confident in their own relationships as a result.

Notably, the same perspective-getting intervention had a negative effect on self-efficacy for students with comparatively higher mentoring relationship satisfaction. Though unexpected, this finding aligns with extant research using perspective focused interventions that has shown these interventions can sometimes lead to detrimental effects (Ku et al., 2015). In the workplace, perspective taking can lead to benefits for a co-worker, but it can also increase self-regulatory resource depletion and hinder the well-being of the individual engaging in perspective taking (Fasbender et al., 2023). More broadly, our findings are consistent with research suggesting that engaging in prosocial behaviors towards others, such as perspective focused or empathic behaviors, can have negative consequences (Gabriel et al., 2018; Koopman et al., 2016; Lin et al., 2022).

One plausible explanation of this cross-over effect is that students with high mentoring satisfaction may have an idealized perception of faculty jobs, which the perspective-getting intervention undermines. This interpretation is consistent with self-discrepancy theory, which posits that individuals experience negative affect when there is misalignment between the attributes they believe they possess (i.e., “the actual self”), the attributes they desire (i.e., “the ideal self”), and the attributes they believe they should possess (i.e., “the ought self”) (Higgins, 1987). Role modeling constitutes a central process in mentoring (Kram, 1988; Lockwood et al., 2002; Scandura, 1992) and doctoral students with highly satisfying mentoring relationships may be modeling their perception of their “ought self” on their faculty mentor. For these students, perspective getting may make them more aware of discrepancies between their idealized perceptions and the actual challenges associated with being a faculty member, thereby undermining their research self-efficacy. Engaging in perspective getting may have inadvertently led to lower self-efficacy for students with higher relationship quality, in part, because they may have been otherwise unaware of their mentors’ perspectives. Students with less satisfying relationships may be less inclined to view their faculty mentors as their “ought self,” and thus less likely to be influenced by gaining insights into the difficulties that faculty mentors’ face.

### 4.3 Contributions and future directions

Our results indicate that perspective-getting interventions are not a panacea for improving doctoral mentoring relationships and that not all students benefit from these programs. However, our intervention did promote a key motivational belief (i.e., self-efficacy) among students who were highly in need of a relational intervention (those with lower mentoring relationship satisfaction). Thus, our results suggest that perspective-getting interventions may be useful when targeted towards specific students. Future research should further assess the moderating effects of relationship satisfaction and utilize more robust measures to understand how particular components of relationship satisfaction (e.g., trust, respect, liking) may influence perspective getting effects. Researchers might also further explore the potential of using perspective getting techniques, such as the ones in this study, to provide targeted supports for students who are less satisfied with the quality of their mentoring relationships.

Another important future direction is trying to broaden the impact of perspective getting messages so that it benefits more students. Our analysis of students’ open-ended text responses indicated that the messages in the perspective getting examples were not accepted or internalized by all students. This was unanticipated, because we conducted extensive pilot testing of our perspective-getting intervention materials during our intervention development process with a goal that materials would resonate with as many students as possible. We purposefully pilot tested with individuals who resembled the target population, and the intervention messages were not critiqued during this testing. Yet, the same intervention materials appeared to elicit negative interactions for a fraction of students at the targeted institution. Potential counterproductive and even back-firing effects have been reported for some motivational interventions in higher education settings (Canning et al., 2019). Findings of our study further highlight the importance of tailoring and precision for intervention messages and underscore the key importance of piloting materials and evaluating students’ responsiveness and internalization of motivational intervention messages when trying to interpret their effects. Future studies should further refine and modify the intervention materials to minimize potential negative reactions prior to further testing, and in turn this may lead to greater benefits for students.

Finally, students in our study participated in two independent intervention sessions: the first session was one week after enrollment in the study and the second session was one month following the first session. We expected that two sessions would be sufficient to change students’ causal attributions and self-efficacy post-intervention. However, we only found moderation effects of perspective getting six months later. These results suggest that getting perspective on negative mentoring experiences may be protective for doctoral students, but such effects take time. Future research should test this possibility more directly by investigating dosage and timing effects of perspective-getting interventions.

### 4.4 Limitations

As with all research, this study has limitations. First, the study was conducted at a single institution with a relatively modest sample of students who were largely homogenous in their demographic characteristics (e.g., mostly women, White, continuing generation). Although our sample is somewhat diverse compared to national enrollment in STEM graduate programs (National Science Foundation, 2023), our sample was underpowered to test if the effects of the intervention were moderated by students’ socio-demographics. An additional limitation is related to measurement. Our self-efficacy scale required substantial modifications to improve measurement model fit. Furthermore, we measured mentoring relationship satisfaction with a single item, which is useful as a broad indicator but precludes a more nuanced understand of potential influences of perspective getting on doctoral student-faculty mentoring relationships.

Finally, we narrowly focused on the potential for perspective getting to influence students’ self-blame and research self-efficacy. Thus, we cannot rule out potential effects of the intervention on other theoretically relevant outcomes, such as increased prosocial behaviors (e.g., helping, sharing, cooperating, empathy) or open mindedness (Ku et al., 2015).

### 4.5 Conclusion

In sum, our study indicates that getting the perspective of their faculty mentors has the potential to preserve the research self-efficacy of doctoral students in less satisfying mentoring relationships, but only months after the intervention. Furthermore, perspective getting has the potential to undermine the research self-efficacy of doctoral students in more satisfying relationships, again only months afterward. These findings provide important insights into the mechanism through which negative interactions with mentors may affect graduate students – specifically as a source of negative social persuasion rather than through their beliefs about the causes of negative interactions. Future research using similarly robust study designs should examine the conditions under which perspective-getting interventions can be effectively implemented to promote STEM doctoral students’ well-being and success.

## Acknowledgements

We thank the members of the Social Psychology of Research Experiences and Education lab and the Biology Education Research Group at the University of Georgia for their feedback that improved our work. This work was supported by the National Institute of General Medical Science of the National Institutes of Health under award number 3T32GM007103-46S1. Additional support was provided by a National Science Foundation Graduate Research Fellowship awarded under grant DGE 1842396 to TTT and the Georgia Athletic Association Professorship of Innovative Science Education. Any opinions, findings, conclusions, or recommendations expressed in this material are those of the authors and do not necessarily reflect the views of any of the funding organizations.

## Author contributions

**Trevor T. Tuma:** Conceptualization, Methodology, Formal analysis, Writing – Original Draft, Visualization. **Heather N. Fedesco:** Conceptualization, Methodology, Investigation, data curation, Writing – Review & Editing, Project administration. **Emily Q. Rosenzweig:** Conceptualization, Methodology, Formal analysis, Writing – Review & Editing, Visualization, Supervision. **Xiao-Yin Chen:** Methodology, Formal analysis, Writing – Review & Editing. **Erin L. Dolan:** Conceptualization, Methodology, Writing – Review & Editing, Supervision, Project administration, Funding acquisition

## REFERENCES

Aikens, M. L., Sadselia, S., Watkins, K., Evans, M., Eby, L., & Dolan, E. L. (2016). A social capital perspective on the mentoring of undergraduate life science researchers: An empirical study of undergraduate–postgraduate–faculty triads. CBE—Life Sciences Education, 15(2), ar16. 10.1187/cbe.15-10-0208

Bandura, A. (1977). Self-efficacy: Toward a unifying theory of behavioral change. Psychological Review, 84(2), 191.

Bandura, A. (1986). Social foundations of thought and action. *Englewood Cliffs*, NJ, 1986(23– 28), 2.

Bandura, A. (1997). Self-Efficacy: The Exercise of Control, Nueva York, NH Freeman.

Barnes, B. J., & Austin, A. E. (2009). The Role of Doctoral Advisors: A Look at Advising from the Advisor’s Perspective. Innovative Higher Education, 33(5), 297–315. 10.1007/s10755-008-9084-x

Blatt, B., LeLacheur, S. F., Galinsky, A. D., Simmens, S. J., & Greenberg, L. (2010). Does perspective-taking increase patient satisfaction in medical encounters? Academic Medicine, 85(9), 1445–1452.

Boese, G. D. B., Stewart, T. L., Perry, R. P., & Hamm, J. M. (2013). Assisting failure-prone individuals to navigate achievement transitions using a cognitive motivation treatment (attributional retraining). Journal of Applied Social Psychology, 43(9), 1946–1955. 10.1111/jasp.12139

Bruneau, E. G., & Saxe, R. (2012). The power of being heard: The benefits of ‘perspective-giving’in the context of intergroup conflict. Journal of Experimental Social Psychology, 48(4), 855–866.

Burt, B. A., McKen, A., Burkhart, J., Hormell, J., & Knight, A. (2019). Black men in engineering graduate education: Experiencing racial microaggressions within the advisor–advisee relationship. Journal of Negro Education, 88(4), 493–508.

Byars-Winston, A., Diestelmann, J., Savoy, J. N., & Hoyt, W. T. (2017). Unique effects and moderators of effects of sources on self-efficacy: A model-based meta-analysis. Journal of Counseling Psychology, 64(6), 645.

Byars-Winston, A., Rogers, J. G., Thayer-Hart, N., Black, S., Branchaw, J., & Pfund, C. (2023). A randomized controlled trial of an intervention to increase cultural diversity awareness of research mentors of undergraduate students. Science Advances, 9(21), eadf9705. 10.1126/sciadv.adf9705

Canning, E. A., Priniski, S. J., & Harackiewicz, J. M. (2019). Unintended consequences of framing a utility-value intervention in two-year colleges. Learning and Instruction, 62, 37– 48.

Chemers, M. M., Zurbriggen, E. L., Syed, M., Goza, B. K., & Bearman, S. (2011). The Role of Efficacy and Identity in Science Career Commitment Among Underrepresented Minority Students. Journal of Social Issues, 67(3), 469–491. 10.1111/j.1540-4560.2011.01710.x

Chen, J. A., & Usher, E. L. (2013). Profiles of the sources of science self-efficacy. Learning and Individual Differences, 24, 11–21. 10.1016/j.lindif.2012.11.002

Choi, B. Y., Park, H., Yang, E., Lee, S. K., Lee, Y., & Lee, S. M. (2012). Understanding Career Decision Self-Efficacy: A Meta-Analytic Approach. Journal of Career Development, 39(5), 443–460. 10.1177/0894845311398042

Clark, R. A., Harden, S. L., & Johnson, W. B. (2000). Mentor relationships in clinical psychology doctoral training: Results of a national survey. Teaching of Psychology, 27(4), 262–268.

Davis, M. H., Conklin, L., Smith, A., & Luce, C. (1996). Effect of perspective taking on the cognitive representation of persons: A merging of self and other. Journal of Personality and Social Psychology, 70(4), 713.

Epley, N., Caruso, E. M., & Bazerman, M. H. (2006). When perspective taking increases taking: Reactive egoism in social interaction. Journal of Personality and Social Psychology, 91(5), 872.

Estrada, M., Hernandez, P. R., & Schultz, P. W. (2018). A Longitudinal Study of How Quality Mentorship and Research Experience Integrate Underrepresented Minorities into STEM Careers. CBE—Life Sciences Education, 17(1), ar9. 10.1187/cbe.17-04-0066

Eyal, T., Steffel, M., & Epley, N. (2018). Perspective mistaking: Accurately understanding the mind of another requires getting perspective, not taking perspective. Journal of Personality and Social Psychology, 114(4), 547.

Fasbender, U., Burmeister, A., & Wang, M. (2020). Motivated to be socially mindful: Explaining age differences in the effect of employees’ contact quality with coworkers on their coworker support. Personnel Psychology, 73(3), 407–430. 10.1111/peps.12359

Fasbender, U., Rivkin, W., & Gerpott, F. H. (2023). Good for you, bad for me? The daily dynamics of perspective taking and well-being in coworker dyads. Journal of Occupational Health Psychology. https://psycnet.apa.org/record/2024-25401-001

Faul, F., Erdfelder, E., Lang, A.-G., & Buchner, A. (2007). G* Power 3: A flexible statistical power analysis program for the social, behavioral, and biomedical sciences. Behavior Research Methods, 39(2), 175–191.

Fedesco, H. N., Kraner, E. R., & Dolan, E. L. (2023). Evaluating the feasibility, utility, and impact of engaging in mentorship assessment to improve doctoral mentoring relationships. New Directions for Teaching and Learning, 2023(176), 95–105. 10.1002/tl.20572

Foundation, N. S. (2023). Diversity and STEM: Women, minorities, and persons with disabilities. NCSES.

Gabriel, A. S., Koopman, J., Rosen, C. C., & Johnson, R. E. (2018). Helping others or helping oneself? An episodic examination of the behavioral consequences of helping at work. Personnel Psychology, 71(1), 85–107. 10.1111/peps.12229

Galinsky, A. D., Ku, G., & Wang, C. S. (2005). Perspective-Taking and Self-Other Overlap: Fostering Social Bonds and Facilitating Social Coordination. Group Processes & Intergroup Relations, 8(2), 109–124. 10.1177/1368430205051060

Galinsky, A. D., Wang, C. S., & Ku, G. (2008). Perspective-takers behave more stereotypically. Journal of Personality and Social Psychology, 95(2), 404.

Gangrade, N., Samuels, C., Attar, H., Schultz, A., Nana, N., Ye, E., & Lambert, W. M. (2024). Mentorship Interventions in Postgraduate Medical and STEM Settings: A Scoping Review. CBE—Life Sciences Education, 23(3), ar33.

Goodyear, R., Crego, C., Johnston, M., & Wph. (1992). Ethical Issues in the Supervision of Student Research: A Study of Critical Incidents. Professional Psychology: Research and Practice, 23, 203–210. 10.1037/0735-7028.23.3.203

Graham, S. (2020). An attributional theory of motivation. Contemporary Educational Psychology, 61, 101861.

Graham, S., & Taylor, A. Z. (2022). The power of asking *why* ?: Attribution retraining programs for the classroom teacher. Theory Into Practice, 61(1), 5–22. 10.1080/00405841.2021.1932160

Grant, A. M., & Berry, J. W. (2011). The Necessity of Others is The Mother of Invention: Intrinsic and Prosocial Motivations, Perspective Taking, and Creativity. Academy of Management Journal, 54(1), 73–96. 10.5465/amj.2011.59215085

Griffin, K. A., Stone, J., Dissassa, D.-T., Hall, T. N., & Clarke, A. (2022). Surviving or flourishing: How relationships with principal investigators influence science graduate students’ wellness. Studies in Graduate and Postdoctoral Education, 14(1), 47–62. 10.1108/SGPE-12-2021-0085

Hamm, J. M., Perry, R. P., Chipperfield, J. G., Hladkyj, S., Parker, P. C., & Weiner, B. (2020). Reframing Achievement Setbacks: A Motivation Intervention to Improve 8-Year Graduation Rates for Students in Science, Technology, Engineering, and Mathematics (STEM) Fields. Psychological Science, 31(6), 623–633. 10.1177/0956797620904451

Hamm, J. M., Perry, R. P., Chipperfield, J. G., Murayama, K., & Weiner, B. (2017). Attribution-based motivation treatment efficacy in an online learning environment for students who differ in cognitive elaboration. Motivation and Emotion, 41(5), 600–616. 10.1007/s11031-017-9632-8

Haynes, T. L., Ruthig, J. C., Perry, R. P., Stupnisky, R. H., & Hall, N. C. (2006). Reducing the Academic Risks of Over-Optimism: The Longitudinal Effects of Attributional Retraining on Cognition and Achievement. Research in Higher Education, 47(7), 755–779. 10.1007/s11162-006-9014-7

Higgins, E. T. (1987). Self-discrepancy: A theory relating self and affect. Psychological Review, 94(3), 319.

Higgins, E. T., & Rholes, W. S. (1978). “Saying is believing”: Effects of message modification on memory and liking for the person described. Journal of Experimental Social Psychology, 14(4), 363–378.

Hoever, I. J., Van Knippenberg, D., Van Ginkel, W. P., & Barkema, H. G. (2012). Fostering team creativity: Perspective taking as key to unlocking diversity’s potential. Journal of Applied Psychology, 97(5), 982.

Hu, L., & Bentler, P. M. (1999). Cutoff criteria for fit indexes in covariance structure analysis: Conventional criteria versus new alternatives. Structural Equation Modeling: A Multidisciplinary Journal, 6(1), 1–55. 10.1080/10705519909540118

Joshi, M., Aikens, M. L., & Dolan, E. L. (2019). Direct Ties to a Faculty Mentor Related to Positive Outcomes for Undergraduate Researchers. BioScience, 69(5), 389–397. 10.1093/biosci/biz039

Kardash, C. M. (2000). Evaluation of undergraduate research experience: Perceptions of undergraduate interns and their faculty mentors. Journal of Educational Psychology, 92(1), 191.

Koopman, J., Lanaj, K., & Scott, B. A. (2016). Integrating the Bright and Dark Sides of OCB: A Daily Investigation of the Benefits and Costs of Helping Others. Academy of Management Journal, 59(2), 414–435. 10.5465/amj.2014.0262

Kram, K. E. (1988). Mentoring at work: Developmental relationships in organizational life. University Press of America. https://psycnet.apa.org/record/1988-97625-000

Ku, G., Wang, C. S., & Galinsky, A. D. (2015). The promise and perversity of perspective-taking in organizations. Research in Organizational Behavior, 35, 79–102.

Lankau, M. J., & Scandura, T. A. (2002). AN INVESTIGATION OF PERSONAL LEARNING IN MENTORING RELATIONSHIPS: CONTENT, ANTECEDENTS, AND CONSEQUENCES. Academy of Management Journal, 45(4), 779–790. 10.2307/3069311

Lazowski, R. A., & Hulleman, C. S. (2016). Motivation Interventions in Education: A Meta-Analytic Review. Review of Educational Research, 86(2), 602–640. 10.3102/0034654315617832

Lee, S. P., McGee, R., Pfund, C., & Branchaw, J. (2015). Mentoring up: Learning to manage your mentoring relationships. The Mentoring Continuum: From Graduate School through Tenure, 133–154.

Lewis, V., Martina, C. A., McDermott, M. P., Chaudron, L., Trief, P. M., LaGuardia, J. G., Sharp, D., Goodman, S. R., Morse, G. D., & Ryan, R. M. (2017). Mentoring Interventions for Underrepresented Scholars in Biomedical and Behavioral Sciences: Effects on Quality of Mentoring Interactions and Discussions. CBE—Life Sciences Education, 16(3), ar44. 10.1187/cbe.16-07-0215

Liang, B., Spencer, R., Brogan, D., & Corral, M. (2008). Mentoring relationships from early adolescence through emerging adulthood: A qualitative analysis. Journal of Vocational Behavior, 72(2), 168–182.

Limeri, L. B., Asif, M. Z., Bridges, B. H. T., Esparza, D., Tuma, T. T., Sanders, D., Morrison, A. J., Rao, P., Harsh, J. A., Maltese, A. V., & Dolan, E. L. (2019). “Where’s My Mentor?!” Characterizing Negative Mentoring Experiences in Undergraduate Life Science Research. CBE—Life Sciences Education, 18(4), ar61. 10.1187/cbe.19-02-0036

Lin, S.-H. J., Poulton, E. C., Tu, M.-H., & Xu, M. (2022). The consequences of empathic concern for the actors themselves: Understanding empathic concern through conservation of resources and work-home resources perspectives. Journal of Applied Psychology, 107(10), 1843.

Lockwood, P., Jordan, C. H., & Kunda, Z. (2002). Motivation by positive or negative role models: Regulatory focus determines who will best inspire us. Journal of Personality and Social Psychology, 83(4), 854.

National Academies of Sciences, Engineering, & Medicine. (2018). Graduate STEM Education for the 21st Century. 10.17226/25038

National Academies of Sciences, Engineering, & Medicine. (2019). The Science of Effective Mentorship in STEMM. 10.17226/25568

National Science Foundation. (2024). NSF’s New Mentoring Requirements for Graduate Students (Proposal and Award Policies and Procedures Guide). https://nsf-gov-resources.nsf.gov/files/nsf24_1.pdf?VersionId=m8jaLa_6ShoF6gQzkyYeMXFMImHgl5jl

Okonofua, J. A., Paunesku, D., & Walton, G. M. (2016). Brief intervention to encourage empathic discipline cuts suspension rates in half among adolescents. Proceedings of the National Academy of Sciences, 113(19), 5221–5226. 10.1073/pnas.1523698113

Paglis, L. L., Green, S. G., & Bauer, T. N. (2006). Does adviser mentoring add value? A longitudinal study of mentoring and doctoral student outcomes. Research in Higher Education, 47(4), 451–476. 10.1007/s11162-005-9003-2

Pajares, F. (1996). Self-Efficacy Beliefs in Academic Settings. Review of Educational Research, 66(4), 543–578. 10.3102/00346543066004543

Parker, P. C., Perry, R. P., Hamm, J. M., Chipperfield, J. G., & Hladkyj, S. (2016). Enhancing the academic success of competitive student athletes using a motivation treatment intervention (Attributional Retraining). Psychology of Sport and Exercise, 26, 113–122. 10.1016/j.psychsport.2016.06.008

Parmar, B. L., Wicks, A. C., & Ginena, K. (2023). The Impact of Employee Stakeholder Orientation on Job Satisfaction and Perspective-Taking. Business & Society, 00076503231182625. 10.1177/00076503231182625

Pengju, W., Xiong, Z., & Hua, Y. (2018). Relationship of mental health, social support, and coping styles among graduate students: Evidence from Chinese universities. Iranian Journal of Public Health, 47(5), 689.

Perry, R. P., & Hamm, J. M. (2017). An attribution perspective on competence and motivation. Handbook of Competence and Motivation: Theory and Application, 2006, 61–84.

Perry, R. P., Stupnisky, R. H., Hall, N. C., Chipperfield, J. G., & Weiner, B. (2010). Bad Starts and Better Finishes: Attributional Retraining and Initial Performance in Competitive Achievement Settings. Journal of Social and Clinical Psychology, 29(6), 668–700. 10.1521/jscp.2010.29.6.668

Peterson, J. L., Bellows, A., & Peterson, S. (2015). Promoting connection: Perspective-taking improves relationship closeness and perceived regard in participants with low implicit self-esteem. Journal of Experimental Social Psychology, 56, 160–164.

Pfund, C., House, S. C., Asquith, P., Fleming, M. F., Buhr, K. A., Burnham, E. L., Gilmore, J. M. E., Huskins, W. C., McGee, R., Schurr, K., Shapiro, E. D., Spencer, K. C., & Sorkness, C. A. (2014). Training Mentors of Clinical and Translational Research Scholars: A Randomized Controlled Trial. Academic Medicine : Journal of the Association of American Medical Colleges, 89(5), 774–782. 10.1097/ACM.0000000000000218

Pfund, C., Spencer, K. C., Asquith, P., House, S. C., Miller, S., & Sorkness, C. A. (2015). Building National Capacity for Research Mentor Training: An Evidence-Based Approach to Training the Trainers. CBE—Life Sciences Education, 14(2), ar24. 10.1187/cbe.14-10-0184

Ragins, B. R. (2011). Relational mentoring: A positive approach to mentoring at work. https://academic.oup.com/edited-volume/28366/chapter/215250104

Ragins, B. R., & Verbos, A. K. (2017). Positive relationships in action: Relational mentoring and mentoring schemas in the workplace. In Exploring positive relationships at work (pp. 91–116). Psychology Press. https://www.taylorfrancis.com/chapters/edit/10.4324/9781315094199-7/positive-relationships-action-belle-rose-ragins-amy-klemm-verbos

Risner, L. E., Morin, X. K., Erenrich, E. S., Clifford, P. S., Franke, J., Hurley, I., & Schwartz, N. B. (2020). Leveraging a collaborative consortium model of mentee/mentor training to foster career progression of underrepresented postdoctoral researchers and promote institutional diversity and inclusion. PloS One, 15(9), e0238518.

Robbins, S. B., Lauver, K., Le, H., Davis, D., Langley, R., & Carlstrom, A. (2004). Do psychosocial and study skill factors predict college outcomes? A meta-analysis. Psychological Bulletin, 130(2), 261.

Roediger, H. L., & Karpicke, J. D. (2006). Test-Enhanced Learning: Taking Memory Tests Improves Long-Term Retention. Psychological Science, 17(3), 249–255. 10.1111/j.1467-9280.2006.01693.x

Rogers, J., Gong, X., Byars-Winston, A., McDaniels, M., Thayer-Hart, N., Cheng, P., Diggs-Andrews, K., Martínez-Hernández, K. J., & Pfund, C. (2022). Comparing the Outcomes of Face-to-Face and Synchronous Online Research Mentor Training Using Propensity Score Matching. CBE—Life Sciences Education, 21(4), ar62. 10.1187/cbe.21-12-0332

Rosenzweig, E. Q., Song, Y., & Clark, S. (2022). Mixed effects of a randomized trial replication study testing a cost-focused motivational intervention. Learning and Instruction, 82, 101660.

Rosenzweig, E. Q., Wigfield, A., & Hulleman, C. S. (2020). More useful or not so bad? Examining the effects of utility value and cost reduction interventions in college physics. Journal of Educational Psychology, 112(1), 166.

Scandura, T. A. (1992). Mentorship and career mobility: An empirical investigation. Journal of Organizational Behavior, 13(2), 169–174. 10.1002/job.4030130206

Sheu, H.-B., Lent, R. W., Miller, M. J., Penn, L. T., Cusick, M. E., & Truong, N. N. (2018). Sources of self-efficacy and outcome expectations in science, technology, engineering, and mathematics domains: A meta-analysis. Journal of Vocational Behavior, 109, 118–136.

Shih, M., Wang, E., Trahan Bucher, A., & Stotzer, R. (2009). Perspective Taking: Reducing Prejudice Towards General Outgroups and Specific Individuals. Group Processes & Intergroup Relations, 12(5), 565–577. 10.1177/1368430209337463

Siegel Robertson, J. (2000). Is attribution training a worthwhile classroom intervention for K–12 students with learning difficulties? Educational Psychology Review, 12, 111–134.

Sitzmann, T., & Yeo, G. (2013). A Meta-Analytic Investigation of the Within-Person Self-Efficacy Domain: Is Self-Efficacy a Product of Past Performance or a Driver of Future Performance? Personnel Psychology, 66(3), 531–568. 10.1111/peps.12035

Sverdlik, A., C. Hall, N., McAlpine, L., & Hubbard, K. (2018). The PhD Experience: A Review of the Factors Influencing Doctoral Students’ Completion, Achievement, and Well-Being. International Journal of Doctoral Studies, 13, 361–388. 10.28945/4113

Tarrant, M., Calitri, R., & Weston, D. (2012). Social Identification Structures the Effects of Perspective Taking. Psychological Science, 23(9), 973–978. 10.1177/0956797612441221

Todd, A. R., & Galinsky, A. D. (2014). Perspective-Taking as a Strategy for Improving Intergroup Relations: Evidence, Mechanisms, and Qualifications. Social and Personality Psychology Compass, 8(7), 374–387. 10.1111/spc3.12116

Tuma, T. T., Adams, J. D., Hultquist, B. C., & Dolan, E. L. (2021). The Dark Side of Development: A Systems Characterization of the Negative Mentoring Experiences of Doctoral Students. CBE—Life Sciences Education, 20(2), ar16. 10.1187/cbe.20-10-0231

Usher, E. L., & Pajares, F. (2008). Sources of Self-Efficacy in School: Critical Review of the Literature and Future Directions. Review of Educational Research, 78(4), 751–796. 10.3102/0034654308321456

Vorauer, J. D., Martens, V., & Sasaki, S. J. (2009). When trying to understand detracts from trying to behave: Effects of perspective taking in intergroup interaction. Journal of Personality and Social Psychology, 96(4), 811.

Vorauer, J. D., & Quesnel, M. (2013). You Don’t Really Love Me, Do You? Negative Effects of Imagine-Other Perspective-Taking on Lower Self-Esteem Individuals’ Relationship Well-Being. Personality and Social Psychology Bulletin, 39(11), 1428–1440. 10.1177/0146167213495282

Walton, G. M., & Cohen, G. L. (2007). A question of belonging: Race, social fit, and achievement. Journal of Personality and Social Psychology, 92(1), 82.

Walton, G. M., & Cohen, G. L. (2011). A Brief Social-Belonging Intervention Improves Academic and Health Outcomes of Minority Students. Science, 331(6023), 1447–1451. 10.1126/science.1198364

Walton, G. M., & Wilson, T. D. (2018). Wise interventions: Psychological remedies for social and personal problems. Psychological Review, 125(5), 617.

Weiner, B. (1985). Attribution Theory. In B. Weiner, Human Motivation (pp. 275–326). Springer New York. 10.1007/978-1-4612-5092-0_7

Yeager, D. S., & Walton, G. M. (2011). Social-psychological interventions in education: They’re not magic. Review of Educational Research, 81(2), 267–301. 10.3102/0034654311405999

Zhang, F., Litson, K., & Feldon, D. F. (2022). Social predictors of doctoral student mental health and well-being. PLOS ONE, 17(9), e0274273. 10.1371/journal.pone.0274273

Zhao, C., Golde, C. M., & McCormick, A. C. (2007). More than a signature: How advisor choice and advisor behaviour affect doctoral student satisfaction. Journal of Further and Higher Education, 31(3), 263–281. 10.1080/03098770701424983

Zhou, H., Majka, E. A., & Epley, N. (2017). Inferring perspective versus getting perspective: underestimating the value of being in another person’s Shoes. Psychological Science, 28(4), 482–493. 10.1177/0956797616687124

