## Supplemental materials for "Seeing isn’t believing? Mixed effects of a perspective-getting intervention to improve mentoring relationships for science doctoral students"

### Table of Contents


| Contents | Pages |
|---|---|
| Full text of *Perspective-getting Condition* & *Control Condition* materials | 2-22 |
| Intervention pilot testing and further refinement | 20 |
| Intervention adherence | 21 |
| Scales used to measure outcome variables | 22-23 |
| Results from measurement model testing and model modifications | 24-26 |
| Concordance with Preregistered Analysis Plan | 27 |
| Examining Pre-Baseline Differences between the *Perspective-getting Condition* & *Control Condition* | 27 |
| Correlations and Descriptive Statistics of Variables | 27 |

### *Perspective-getting Condition* Materials

**Session 1**

Welcome! Thanks again for agreeing to participate in this study. We appreciate your continued participation in our study about improving doctoral students' experiences in their life science programs. To remind you, the purpose of the study activities is to gather your feedback regarding materials we have created for graduate students.

Mentoring students in research is a big part of being a faculty member. Graduate students often learn about what it's like to be a mentor by observing how other faculty – usually their research advisors – mentor graduate students. In order to expose graduate students to different mentoring perspectives so they can continue to think about what it might be like to be a mentor in the future, we want to provide you with faculty members' perspectives about mentorship.

In this module, we will share with you some of the benefits and challenges that principal investigator (PIs) experience when mentoring graduate students. Then, we will ask for your feedback so we can better understand how graduate students might use this information. We expect that this will take you ~15-30 minutes.

Thanks in advance for your time.

To gain a deeper understanding of the rewards and challenges of mentoring graduate students, we asked PIs to tell us more about their experiences of being a mentor.

> "As someone with a family, being a PI is great because I can set my own calendar to make sure I leave in time to pick up my kids from daycare, or I can make time to attend school functions. But, when I had kids, I realized early on that I needed to tightly schedule my time to ensure I am spending time with my family while also being a good mentor. Sometimes, in order to protect my time with my family, I need to set boundaries around my availability for the people in my lab. This means I often have to tell my graduate students that I can't review their work last minute or answer their questions over the weekend. It has nothing to do with the students, I always want to help them whenever they need me but, when I became a parent, I just had less time to give." – Assistant Professor

> "I care a lot about making sure students feel like their contributions matter in my lab. But as a PI, when there are disagreements among the group about the best course of action, I usually have to make the call. Sometimes if we go in one direction over another, I worry that students will take it personally, as if I'm saying their ideas aren't good enough. However, that's just not true – the discussions and suggestions all contribute to the problem-solving process. I am trying to work on helping my students understand that I value all of their input, and if the group doesn't end up taking their suggestions it doesn't have anything to do with their skills as a researcher." – Professor

“One of the joys of the job for me is working with grad students in my lab. When I was a grad student, my PI was a great mentor and I want to pay it forward to help a new generation of scientists in training. I really enjoy helping students, but it takes time. And sometimes I’m not able to provide the right support and guidance that students need. I plan to do a better job of fostering a network of mentors for my students so they know there are other people they can turn to for support if I happen to be unavailable or perhaps not the best person to help in the particular situation.” – Associate Professor

“One challenge I experience as a PI is that when I'm working on grants, it takes away time I could spend mentoring people in my group. I need to do it because that's how I can keep the team going. I’m sure my focus on getting funding can seem like I am not invested in my team’s current work. But if I don’t have enough funding coming in, not only does the research stall, but I will need to quickly find other ways to fund people in my group. It’s a lot of work but I find that writing proposals can be really fun because it involves telling a scientific story and convincing reviewers that the research you are proposing is worth taxpayer investment. And it gives me lots of opportunities to mentor students in my lab on the process of applying for grants.” – Assistant Professor

We’d like to know what you think of each example. Please answer the following questions about each faculty quote.

- How informative is this quote to you?
- How relevant is this quote to your own experiences as a graduate student?

Quote 1

“As someone with a family, being a PI is great because I can set my own calendar to make sure I leave in time to pick up my kids from daycare, or I can make time to attend school functions. But, when I had kids, I realized early on that I needed to tightly schedule my time to ensure I am spending time with my family while also being a good mentor. Sometimes, in order to protect my time with my family, I need to set boundaries around my availability for the people in my lab. This means I often have to tell my graduate students that I can’t review their work last minute or answer their questions over the weekend. It has nothing to do with the students, I always want to help them whenever they need me but, when I became a parent, I just had less time to give.” – Assistant Professor

We’d like to know what you think of each example. Please answer the following questions about each faculty quote.

- How informative is this quote to you?
- How relevant is this quote to your own experiences as a graduate student?

Quote 2

“I care a lot about making sure students feel like their contributions matter in my lab. But as a PI, when there are disagreements among the group about the best course of action, I usually have to make the call. Sometimes if we go in one direction over another, I worry that students will take it personally, as if I’m saying their ideas aren’t good enough. However, that’s just not true – the discussions and suggestions all contribute to the problem-solving process. I am trying to work on helping my students understand that I value all of their input, and if the group doesn't end up taking their suggestions it doesn't have anything to do with their skills as a researcher." – Professor

We'd like to know what you think of each example. Please answer the following questions about each faculty quote.

- How informative is this quote to you?
- How relevant is this quote to your own experiences as a graduate student?

<u>Quote 3</u>
"One of the joys of the job for me is working with grad students in my lab. When I was a grad student, my PI was a great mentor and I want to pay it forward to help a new generation of scientists in training. I really enjoy helping students, but it takes time. And sometimes I'm not able to provide the right support and guidance that students need. I plan to do a better job of fostering a network of mentors for my students so they know there are other people they can turn to for support if I happen to be unavailable or perhaps not the best person to help in the particular situation." – Associate Professor

We'd like to know what you think of each example. Please answer the following questions about each faculty quote.

- How informative is this quote to you?
- How relevant is this quote to your own experiences as a graduate student?

<u>Quote 4</u>
"One challenge I experience as a PI is that when I'm working on grants, it takes away time I could spend mentoring people in my group. I need to do it because that's how I can keep the team going. I'm sure my focus on getting funding can seem like I am not invested in my team's current work. But if I don't have enough funding coming in, not only does the research stall, but I will need to quickly find other ways to fund people in my group. It's a lot of work but I find that writing proposals can be really fun because it involves telling a scientific story and convincing reviewers that the research you are proposing is worth taxpayer investment. And it gives me lots of opportunities to mentor students in my lab on the process of applying for grants." – Assistant Professor

Now we would like you to rank the quotes in order of your most to least favorite quotes. Drag the quotes shown below into the order that you prefer. The **<u>top quote</u> should be your <u>favorite</u>** quote, and the **<u>bottom quote</u> should be your <u>least</u> favorite** quote.

> "As someone with a family, being a PI is great because I can set my own calendar to make sure I leave in time to pick up my kids from daycare, or I can make time to attend school functions. But, when I had kids, I realized early on that I needed to tightly schedule my time to ensure I am spending time with my family while also being a good mentor. Sometimes, in order to protect my time with my family, I need to set boundaries around my availability for the people in my lab. This means I often have to tell my graduate students that I can't review their work last minute or answer their questions over the weekend. It has nothing to do with the students, I always want to help them whenever

they need me but, when I became a parent, I just had less time to give." – Assistant Professor

"I care a lot about making sure students feel like their contributions matter in my lab. But as a PI, when there are disagreements among the group about the best course of action, I usually have to make the call. Sometimes if we go in one direction over another, I worry that students will take it personally, as if I'm saying their ideas aren't good enough. However, that's just not true – the discussions and suggestions all contribute to the problem-solving process. I am trying to work on helping my students understand that I value all of their input, and if the group doesn't end up taking their suggestions it doesn't have anything to do with their skills as a researcher." – Professor

"One of the joys of the job for me is working with grad students in my lab. When I was a grad student, my PI was a great mentor and I want to pay it forward to help a new generation of scientists in training. I really enjoy helping students, but it takes time. And sometimes I'm not able to provide the right support and guidance that students need. I plan to do a better job of fostering a network of mentors for my students so they know there are other people they can turn to for support if I happen to be unavailable or perhaps not the best person to help in the particular situation." – Associate Professor

"One challenge I experience as a PI is that when I'm working on grants, it takes away time I could spend mentoring people in my group. I need to do it because that's how I can keep the team going. I'm sure my focus on getting funding can seem like I am not invested in my team's current work. But if I don't have enough funding coming in, not only does the research stall, but I will need to quickly find other ways to fund people in my group. It's a lot of work but I find that writing proposals can be really fun because it involves telling a scientific story and convincing reviewers that the research you are proposing is worth taxpayer investment. And it gives me lots of opportunities to mentor students in my lab on the process of applying for grants." – Assistant Professor

What about your top-ranked quote caused you to rank it first? *Write in response.*

Now we would like your feedback. On the next page, in 1-2 paragraphs, please write about ways in which the information you read informs how you think about your interactions and challenges with your own research advisor. Please also write about how this information might influence your level of interest in being a research mentor.

To help you reflect on this prompt, read and review the following example responses from graduate students who read these materials previously:

<u>Quote 1:</u>
"It was reassuring to read more about the challenges PIs have of making decisions on projects, especially when people in the lab have different ideas. In my lab, we have weekly meetings where people give updates on their individual projects and turn to the group for ideas on how to improve what they are working on. I can definitely relate to feeling like my ideas aren't good enough when we are in a lab meeting and the group doesn't take my suggestions. I'm going to

try to remember that during those meetings, everyone is trying to help each other make the lab's projects as successful as possible. And even if we go in a different direction, that my contributions help the team get closer to coming up with the best course of action as part of the brainstorming process. All this information confirmed my desire to be a PI because I think I have the skillset to be an effective research mentor and I want to help future generations of scientists." – 2nd year graduate student

Quote 2:
"It can be hard to find a time to meet with my advisor and they sometimes cancel our one-on-one meetings, which is really frustrating. But it's helpful to realize just how much advisors have to deal with, that they are pulled in many different directions, and that I shouldn't take cancelled meetings personally. Next time my advisor cancels a meeting, I'll ask if there is a way for me to get feedback that might be easier to fit into their schedule. Maybe I can email a brief explanation of the results I want to share and the questions I want to ask, so they can give feedback at a time that works for them. I also appreciated knowing just how flexible PIs can be with their schedules and reading about the importance of setting boundaries to maintain a work/life balance. I'm still not sure whether I want to be a PI but this definitely feels like a perk." – 3rd year graduate student

Quote 3:
"This information helped me think differently about the times when I feel like I don't know how to move forward with my research because my advisor isn't always around for me to ask questions. I usually have to email them when I run into issues but in many cases, it would be so much easier if I could just pop into their office real quick. Sometimes I feel like my advisor cares about other more interesting projects because I don't often see them in the lab. But I remembered that my advisor has to take care of their aging parents who aren't doing that well. My advisor's a pretty private person so it was useful for me to remind myself that they've got a life outside the lab. Those obligations are likely pretty stressful and they are probably doing the best they can. I want people to understand that I have other things going on besides grad school so I should probably be more understanding myself. This information did not change my interest in being a research mentor, but if this is part of my future career, it helped me think about how I should be more open with the people I'm mentoring about my schedule." – 4th year graduate student

Please write your response below. *Write in response*

**Session 2**

Welcome! Thanks again for agreeing to participate in this study. We appreciate your continued participation in our study about improving doctoral students' experiences in their life science programs. To remind you, the purpose of the study activities is to gather your feedback regarding materials we have created for graduate students.

A few weeks ago, you read about the rewards and challenges of mentoring graduate students and you wrote about how you would apply what you learned. Write down anything you remember from that material. *Write in response.*

Today, we would like you to continue thinking about what it's like to mentor graduate students by reading some more examples about the experiences of faculty members. Just like last time, we would like your feedback on these examples. This will take 15-30 minutes.

When you are ready to read the examples, click the arrow.

To gain a deeper understanding of the rewards and challenges of mentoring graduate students, we asked PIs to tell us more about their experiences of being a mentor. Please read the following examples. Then, we will ask you a few questions about each one.

> "I really enjoy working with multiple students in my lab because it means we have a lot of minds working on our research questions and my grad students make such great contributions to our projects. But the more students you have, the more challenges there can be. I've done some mentoring professional development and agree with the suggestions made in those workshops, it's just hard to do them all. For example, I know I am sometimes not as available as I'd like to be because we have a lot of projects going on, plus I have other responsibilities as a faculty member. What I have taken away from some of those mentoring workshops is just being more transparent with my students when some weeks get super busy. I don't want to make it seem like I'm making excuses or that I can't get all the work done, but ultimately, I think it may help my students better understand why some weeks can be harder." – Assistant Professor

> "One of the challenges I face as a research mentor is finding the right balance between having students take ownership of their projects but also making sure they are set up for success. I know there's a lot to learn when the research doesn't go the way you had hoped, but I also want to make sure certain issues don't delay a student's ability to graduate on time, make them less competitive for awards or grants, or make them less marketable. I'm sure students sometimes feel like I don't trust their instincts when I disagree with their ideas, and I certainly don't want them to feel that way. In these moments, I should probably do a better job of explaining what I see are the bigger picture risks of taking certain paths." – Associate Professor

> "In my lab, I work with a lot of really creative and hardworking people. We all have similar research interests, but because students are carving out different projects for their theses and dissertations, I find that I am more knowledgeable about some projects than

others. Because of this, during our lab meetings I work hard to make sure I am showing equal attention to everyone's projects and contributions. The last thing I want is for any of my students to think I don't like them as much as others or to question whether they are good enough to be in my lab because, the fact is, I think all of my students are capable and have the potential to be great researchers."– Professor

Just like last time, we'd like to know what you think of each example. Please answer the following questions about each faculty quote.

- How informative is this quote to you?
- How relevant is this quote to your own experiences as a graduate student?

Quote 1

"I really enjoy working with multiple students in my lab because it means we have a lot of minds working on our research questions and my grad students make such great contributions to our projects. But the more students you have, the more challenges there can be. I've done some mentoring professional development and agree with the suggestions made in those workshops, it's just hard to do them all. For example, I know I am sometimes not as available as I'd like to be because we have a lot of projects going on, plus I have other responsibilities as a faculty member. What I have taken away from some of those mentoring workshops is just being more transparent with my students when some weeks get super busy. I don't want to make it seem like I'm making excuses or that I can't get all the work done, but ultimately, I think it may help my students better understand why some weeks can be harder." – Assistant Professor

We'd like to know what you think of each example. Please answer the following questions about each faculty quote.

- How informative is this quote to you?
- How relevant is this quote to your own experiences as a graduate student?

Quote 2

"One of the challenges I face as a research mentor is finding the right balance between having students take ownership of their projects but also making sure they are set up for success. I know there's a lot to learn when the research doesn't go the way you had hoped, but I also want to make sure certain issues don't delay a student's ability to graduate on time, make them less competitive for awards or grants, or make them less marketable. I'm sure students sometimes feel like I don't trust their instincts when I disagree with their ideas, and I certainly don't want them to feel that way. In these moments, I should probably do a better job of explaining what I see are the bigger picture risks of taking certain paths." – Associate Professor

We'd like to know what you think of each example. Please answer the following questions about each faculty quote.

- How informative is this quote to you?
- How relevant is this quote to your own experiences as a graduate student?

Quote 3

"In my lab, I work with a lot of really creative and hardworking people. We all have similar research interests, but because students are carving out different projects for their theses and dissertations, I find that I am more knowledgeable about some projects than others. Because of this, during our lab meetings I work hard to make sure I am showing equal attention to everyone’s projects and contributions. The last thing I want is for any of my students to think I don’t like them as much as others or to question whether they are good enough to be in my lab because, the fact is, I think all of my students are capable and have the potential to be great researchers.”– Professor

Now we would like you to rank the quotes in order of your most to least favorite quotes. Drag the quotes shown below into the order that you prefer. The **top quote should be your favorite** quote, and the **bottom quote should be your least favorite** quote.

> “I really enjoy working with multiple students in my lab because it means we have a lot of minds working on our research questions and my grad students make such great contributions to our projects. But the more students you have, the more challenges there can be. I’ve done some mentoring professional development and agree with the suggestions made in those workshops, it's just hard to do them all. For example, I know I am sometimes not as available as I’d like to be because we have a lot of projects going on, plus I have other responsibilities as a faculty member. What I have taken away from some of those mentoring workshops is just being more transparent with my students when some weeks get super busy. I don’t want to make it seem like I’m making excuses or that I can’t get all the work done, but ultimately, I think it may help my students better understand why some weeks can be harder.” – Assistant Professor

> “One of the challenges I face as a research mentor is finding the right balance between having students take ownership of their projects but also making sure they are set up for success. I know there’s a lot to learn when the research doesn’t go the way you had hoped, but I also want to make sure certain issues don’t delay a student’s ability to graduate on time, make them less competitive for awards or grants, or make them less marketable. I’m sure students sometimes feel like I don’t trust their instincts when I disagree with their ideas, and I certainly don’t want them to feel that way. In these moments, I should probably do a better job of explaining what I see are the bigger picture risks of taking certain paths.” – Associate Professor

> “In my lab, I work with a lot of really creative and hardworking people. We all have similar research interests, but because students are carving out different projects for their theses and dissertations, I find that I am more knowledgeable about some projects than others. Because of this, during our lab meetings I work hard to make sure I am showing equal attention to everyone’s projects and contributions. The last thing I want is for any of my students to think I don’t like them as much as others or to question whether they are good enough to be in my lab because, the fact is, I think all of my students are capable and have the potential to be great researchers.”– Professor

What about your top-ranked quote caused you to rank it first? *Write in response*

During the last session, we asked you to write about how seeing PIs' perspectives about mentorship can positively impact your experience in your doctoral program. Today we would like you to write about the same topic again.

We want your feedback a second time because you now have read more examples of the rewards and challenges of being a research mentor. We also would like your feedback now that some time has passed since reviewing the module.

On the next page, in 1-2 paragraphs, please write about ways in which the information you read informs how you think about your interactions and challenges with your own research advisor. Please also write about how this information might influence your level of interest in being a research mentor. You can write about the same things you reflected on last time or something else.

To help you reflect on this prompt, read and review the following example responses from graduate students who read these materials previously. You have seen these examples before but we are showing them again in case you would like a reminder:

> "It was reassuring to read more about the challenges PIs have of making decisions on projects, especially when people in the lab have different ideas. In my lab, we have weekly meetings where people give updates on their individual projects and turn to the group for ideas on how to improve what they are working on. I can definitely relate to feeling like my ideas aren't good enough when we are in a lab meeting and the group doesn't take my suggestions. I'm going to try to remember that during those meetings, everyone is trying to help each other make the lab's projects as successful as possible. And even if we go in a different direction, that my contributions help the team get closer to coming up with the best course of action as part of the brainstorming process. All this information confirmed my desire to be a PI because I think I have the skillset to be an effective research mentor and I want to help future generations of scientists." – 2nd year graduate student

> "It can be hard to find a time to meet with my advisor and they sometimes cancel our one-on-one meetings, which is really frustrating. But it's helpful to realize just how much advisors have to deal with, that they are pulled in many different directions, and that I shouldn't take cancelled meetings personally. Next time my advisor cancels a meeting, I'll ask if there is a way for me to get feedback that might be easier to fit into their schedule. Maybe I can email a brief explanation of the results I want to share and the questions I want to ask, so they can give feedback at a time that works for them. I also appreciated knowing just how flexible PIs can be with their schedules and reading about the importance of setting boundaries to maintain a work/life balance. I'm still not sure whether I want to be a PI but this definitely feels like a perk." – 3rd year graduate student

> "This information helped me think differently about the times when I feel like I don't know how to move forward with my research because my advisor isn't always around for me to ask questions. I usually have to email them when I run into issues but in many cases, it would be so much easier if I could just pop into their office real quick.

Sometimes I feel like my advisor cares about other more interesting projects because I don’t often see them in the lab. But I remembered that my advisor has to take care of their aging parents who aren’t doing that well. My advisor’s a pretty private person so it was useful for me to remind myself that they’ve got a life outside the lab. Those obligations are likely pretty stressful and they are probably doing the best they can. I want people to understand that I have other things going on besides grad school so I should probably be more understanding myself. This information did not change my interest in being a research mentor, but if this is part of my future career, it helped me think about how I should be more open with the people I’m mentoring about my schedule.” – 4th year graduate student

Please write your response below *Write in response*

### *Control Condition* Materials

#### Session 1

Welcome! Thanks again for agreeing to participate in this study. We appreciate your continued participation in our study about improving doctoral students' experiences in their life science programs. To remind you, the purpose of the study activities is to gather your feedback regarding materials we have created for graduate students.

One important career development skill for graduate students is being an effective research mentor because it is applicable to many career paths a student might pursue. We want to help graduate students develop these skills by sharing some strategies for mentoring others in research.

In this module, we will share tips for establishing successful mentoring relationships with researchers-in-training. Then, we will ask for your feedback so we can better understand how graduate students might use this information. We expect that this will take you ~15-30 minutes.

Thanks in advance for your time.

Regardless of whether graduate students pursue careers within or outside of academia, if they continue to conduct research, they may find themselves mentoring newer scientists who are learning how to carry out research. In fact, many academic and non-academic job interviews will ask candidates about their research mentoring experiences and philosophies.

To prepare for these experiences, it is important for graduate students to be aware of strategies to effectively mentor others in research. Learning about these approaches can help graduate students improve their mentorship of current undergraduate researchers and ensure that they are well-prepared to succeed in future positions that require them to mentor others.

Here, we will highlight some strategies for mentoring undergraduate researchers that were developed by the Center for the Improvement of Mentored Experience in Research at the University of Wisconsin. While these tips focus on mentoring undergraduate students, they are applicable to other mentoring relationships and can help ensure a successful experience for both mentees and mentors.

**Orienting New Mentees**
Initial training is an important part of the process of mentoring a new mentee. The goal is to equip mentees with knowledge and resources to help them get started in their research. Below is a to-do list you can follow when helping a new undergraduate researcher get oriented:

- Outline expectations and guidelines for lab work (e.g., expected work time, how to keep a lab notebook, expected meeting times and frequency, etc.).
- Review required training and safety procedures (e.g., hazardous waste safety, animal care, human subjects, etc.).
- Show students how to find and use lab protocols, data or code repositories, or other informational tools commonly used in the lab.

- Show students how to use common equipment.
- Share key papers and show how to search relevant literature.

**Getting to Know Each Other and Setting Mutually Agreeable Expectations**
Mentors and mentees should make an effort to get to know each other at the start of their relationship. This is a good time to begin defining your working relationship and to establish expectations. Here are some questions to get you started:

- Possible questions to ask mentees (you do not need to ask all of them):
    - Tell me about yourself. What year in school are you? Where do you call home?
    - What is your major? What science topics are interesting to you?
    - What career(s) are you thinking of pursuing?
    - Tell me about why you want to do research. How will doing research help you reach your career goals?
    - What are you hoping to get out of this research experience? What would a successful research experience look like for you?
    - Do you have any previous research experience? If so, what did you do? What did you like about it? What did you dislike about it?
    - What are you looking for from a research mentor?
    - Tell me about your workstyle. What time of day do you work best?

- Possible talking points for mentors (you do not need to talk about all of them):
    - Introduce yourself and your scientific/educational journey so far. Share where you are in your program/career. Share where you grew up/where you have lived/where you call home.
    - Share what you hope to get out of the experience of working with an undergraduate researcher.
    - Explain what success in this research would look like to you. What skills (e.g., technical, communication) are you aiming for your mentee to develop?
    - Explain the proper channels of communication (e.g., email, slack, texting).
    - Clearly state how many hours per week you expect your mentee to work in the lab. Are there specific times of day that you expect your student to be in the lab? How should time be tracked, if at all? What should your mentee do if they are ill or have other conflicts (e.g., several exams in a single week, doctor's appointment) that may require missing their scheduled lab time?

**Considerations for Selecting Research Projects**
The type of project that an undergraduate researcher works on can substantially influence the quality of their research experience. It is important to select a project that is both feasible and meaningful. Consider the following questions when choosing a project for undergraduate researchers:

- Does the project have a reasonable scope so that the student can make some progress and achieve some milestones given how long they will be involved in the lab?
- Does the student have the knowledge and skills to carry out the project? If not, how will they be supported in developing the requisite knowledge and skills?
- Is the project sufficiently challenging so that it will be interesting and motivating to work on? How can the project be tweaked so it is the right level of challenge? Are there ways

to incorporate learning a few techniques or types of research tasks to keep things interesting, but not so many that it becomes overwhelming?
- How is the student's project connected to the ongoing work in the lab and/or other work in the field? How will you help the student understand these connections?
- Can the student have a choice between a few project ideas that they would like to pursue?
- What actions will be taken if the initial project stalls or turns out to not be feasible? Will there be other projects the student can work on?

**Helping Mentees Prepare to Present Their Work**
Mentees are often expected to present their work in a talk, poster, or paper. They should start preparing early and think about how to share their research progress and results in ways that are understandable and interesting to the target audience(s). Here are points for mentors and mentees to consider when planning presentations:
- Focus on the big picture. What do you and your mentee want people to remember about the research? Why should the audience care?
- If your mentee is creating a poster, show them a few examples of posters you think are more or less effective. Discuss the strengths and limitations of different posters with your mentee.
- Ask your mentee to think about the audience for their presentation. What do they think the audience would like to hear or see?
- Decide together on the starting material. Will the student have access to your text, data, figures, etc., or are they building their own from scratch? What would you both prefer? What is reasonable given the timeline?
- Show them how to create a poster or presentation using the technology that is available. Make sure they are aware of any templates, guidelines, or restrictions (e.g., dimensions of a poster, duration of a presentation).
- Have your student practice their poster or oral presentation more than once. For instance, they can practice for other members of your lab or department, or even for their roommates or friends.

We'd like to know what you think of this module. Please rate the degree to which you agree with the following statements (*1 = strongly disagree – 5 = strongly agree)*:

1. *I like this module.*
2. *This module was informative to me.*
3. *This module was relevant to my own experiences as a graduate student.*

Please identify the section that was most helpful to you:
- Orienting New Mentees
- Getting to Know Each Other and Setting Mutually Agreeable Expectations
- Considerations for Selecting Research Projects
- Helping Mentees Prepare to Present Their Work
- Prefer not to respond

Now we would like your feedback. On the next page, in 1-2 paragraphs, please write about ways that knowing more about effective strategies for mentoring undergraduate researchers can positively impact your experience in your doctoral program and/or mentoring experiences you might have in your future career.

To help you reflect on this prompt, you can read and review the following example responses from graduate students who read these materials previously:

<u>Quote 1:</u>
"As I think about how I have mentored undergraduate students during my Ph.D., some of the problems I've encountered have been because I assume students know more about how to do lab work than they actually do. I liked the tips about getting a better sense of students' research experience prior to starting in the lab so I know what they are and aren't familiar with. And then I should definitely spend more time walking through some basics like lab protocols, showing where to find useful information, giving them a refresher on how to use certain equipment, and providing tips for creating figures when presenting their results. Doing this at the outset will probably reduce some frustration that can arise later on." - 3rd year graduate student

<u>Quote 2:</u>
"We have a bunch of undergraduate researchers in our lab, often working on projects for course credit. I have mentored a few of them. This module reminded me of the problems that arose when my PI and I hadn't thought in advance about the scope of their projects. Some undergrads have worked on projects that were probably too simple, or where they did one thing for the entire project. It was helpful to read the tips on what makes a suitable project and that it should have some challenging elements and involve different tasks. In the future I'll be more mindful to help shape or select projects that include some variety for students." - 4th year graduate student

<u>Quote 3:</u>
"In the past, I usually let students come up with a first draft of a presentation on their own. Then, during a meeting, I have them present what they came up with and I give them a lot of feedback afterwards. I can see how reviewing some of the guidelines for designing a research presentation before they start working on it would be more helpful and might cut down on the amount of editing they have to do after I see the presentation for the first time." - 4th year graduate student

Please write your response below: *write in response*

**Session 2**

A few weeks ago, you read about strategies for mentoring others in research and you wrote about how you would apply what you learned. Write down anything you remember from that material. *Write in response*

Today, we would like to share more strategies for developing and sustaining successful research mentoring relationships, which can be applicable to many career paths a graduate student might pursue. Just like last time, we would like your feedback on this material. This will take ~15-30 minutes.

When you are ready to read the strategies, click the arrow.

Last time you read about a variety of strategies for mentoring newer researchers to ensure a successful experience for both the mentees and mentors. We want to share more tips that can improve your current experiences mentoring undergraduate researchers and help you prepare to succeed in future positions that require you to mentor others.)

**Checking on Progress and Revisiting Expectations**
After mentors and mentees have spent some time getting to know each other and working together, it is important to check in periodically about how things are going and revisit expectations. This is also a good time to ensure that the goals and tasks of the research project are clear and to establish or refine a timeline for completion of specific tasks.

- Possible questions to ask mentees (you do not need to ask all of them):
    - What do you like best about working in the lab so far? What aspects of your experience are going well?
    - What do you find most challenging? How can I help you with this?
    - What have you learned about working in a lab that is a surprise to you?
    - Now that you have had some time to work on the research, what questions do you have about the research? What aspects are the most exciting and interesting? What aspects are less interesting?
    - Which techniques, tasks, or protocols do you find most challenging? How can I help you master them?
    - Are we meeting for the right amount of time? Do you feel like we should meet more or less?
    - What feedback do you have about my mentorship? What should I be sure to keep doing because it is working for you? What are 1-3 things you would like me to do differently to make your experience better?
- Possible talking points for mentors (you do not need to talk about all of them):
    - Describe what you see as your mentee's greatest strength(s) in the lab and in doing research so far.
    - Share your thoughts on 1-3 areas you'd like to see your mentee focus on developing. How do you suggest they do this? How can you facilitate this process?

    - Describe the next steps, immediate milestones, and anticipated products (e.g., data, figures, refined protocols, etc.) for the mentee's research as well as the longer-term plan for the research.
    - Share your estimate of the time you anticipate the project and/or specific research tasks will take, and check with your mentee to see if they think this is reasonable.
    - Share what is working well about your interactions with your mentee from your perspective. Consider sharing one or two ways you think your interactions could be improved.

**Tips for Sending/Receiving Feedback**
Mentoring is a skill that develops over time and can always be improved. In addition, each mentoring relationship is unique – what may work for one mentee may not work for another. As you check on your mentee's progress and revisit expectations, you are opening up dialogue for you both to send and receive feedback. To ensure those conversations are successful, consider the following tips:

- Decide what mode of communication would be best to solicit feedback on how the mentoring relationship is going (e.g., in-person conversation, written feedback with a follow-up conversation, etc.)
- When gathering feedback, start by reviewing the specific aspects of your relationship that are going well. Acknowledging positives has multiple benefits, including lowering levels of stress and making it easier to work through problems and concerns.
- When identifying areas that need improvement, it can be easy to jump to defending against or disagreeing with feedback. Instead, consider these steps:
    - Make sure you understand the feedback. Repeat back the feedback in your own words and check whether your summaries are accurate. Ask questions to clarify your understanding, including asking for examples to illustrate.
    - If you find yourself disagreeing with the feedback even after you are sure you understand it, attempt to figure out why you see things differently. Consider what is fair or reasonable about their feedback and what makes sense about what they are saying.
- Then consider what needs to be done next to act on the feedback you received:
    - What feedback seems most important to act on? Can you agree on what should be prioritized?
    - If you have already tried to act on some of the feedback, assess whether that was effective or not and discuss why. Can you try a similar approach but with some modifications?
    - How can you build on this momentum to continue to improve your working relationship? For instance, how can you create the time and space to continue checking in on how things are going? Think about and discuss how often you should check in and what process you will put in place to ensure you both follow through.

**Strategies for Ensuring a Strong Finish**
It's not always clear how to wrap up time with a mentee or conclude your work together. To ensure a strong finish for mentees, consider the following questions:

- Possible questions to ask mentees (you do not need to ask all of them):

- Do you feel that you achieved the goals you outlined at the beginning? Why or why not?
- What aspects of your research and progress are you most proud of? What aspects of your research and progress have been most frustrating for you? Is there anything we could have done differently to help you navigate those frustrations?
- What, if anything, would you still like to accomplish with regard to your research project?
- How would you like to maintain contact with your mentor once the program has ended?

- Possible talking points for mentors (you do not need to talk about all of them):
  - What do you think has been your mentee's greatest accomplishment in their research and/or development?
  - What have you learned from mentoring this student in research? How have you benefited from your work together?
  - How do you feel about the progress you and your mentee have made on this research project?
  - What do you see as next steps for the research project? Do you envision your mentee being involved? If so, in what ways and to what extent?
  - Would you like to maintain contact with your mentee once they conclude their time in the lab? If so, how would you like to go about this (e.g., check in by email or text every 6-12 months, plan to meet online/in-person if/when the research is at the writing stage, meet more informally at a conference)?

We'd like to know what you think of this module. Please rate the degree to which you agree with the following statements (*1 = strongly disagree – 5 = strongly agree)*:

1. *I like this module.*
2. *This module was informative to me.*
3. *This module was relevant to my own experiences as a graduate student.*

Now we would like you to identify the section that was the most helpful to you:

- Checking on Progress and Revisiting Expectations
- Tips for Sending/Receiving Feedback
- Strategies for Ensuring a Strong Finish
- Prefer not to respond

During the last session, we asked you to write about how reading about strategies for mentoring undergraduate researchers can positively impact your experience in your doctoral program. Today we would like you to write about the same topic again.

We want your feedback a second time because you have read more mentoring tips. We also would like your feedback now that some time has passed since reviewing the first module.

On the next page, in 1-2 paragraphs, please write about ways in which the information you read can positively impact your experience in your doctoral program and/or mentoring experiences you might have in your future career. You can write about the same things you reflected on last time or something else.

To help you reflect on this prompt, read and review the following example responses from graduate students who read these materials previously:

> "I really liked the tips for checking in with students and making sure I revisit expectations. It was also helpful to think about how best to receive feedback, as I know I sometimes get defensive when I receive criticism, and I'm sure the undergrads I've worked with have too. I expect that I will mentor newer scientists in my future career so keeping these principles in mind will be helpful." - 3rd year graduate student
>
> "I appreciated the questions about asking mentees what's working well, what they like about working in the lab, what surprised them, and so on. These feel like questions that would allow me to continue building a good connection with the students in the lab while also still being really relevant to their work." - 2nd year graduate student
>
> "I never really know how best to end a relationship with an undergrad working in the lab. I feel like I just sort of say, 'see ya later!' and that's it. I can see how asking some of the questions about maintaining contact afterwards and how best to facilitate that would make things less awkward and just a better way to wrap things up. My goal is to work in industry where I'll probably work with a lot of interns over the years so this seems really useful." - 4th year graduate student

Please write your response below: *Write in response.*

**Intervention pilot testing and further refinement**

After drafting the *Perspective-getting Condition* materials, we refined them through a series of 1-hour focus groups with life science doctoral students, postdoctoral researchers, and faculty who had lived experiences as mentees or mentors (*n* = 15 total across three, five-person groups). These individuals were asked to examine the *Perspective-getting Condition* materials to identify parts they found most and least appealing, and to identify any language that they found to be off-putting, irrelevant, or unpleasant. We used the focus group feedback to edit the materials, particularly the quotations, to be more authentic and reflective of doctoral student-faculty mentoring relationships.

We sought another round of feedback on the revised *Perspective-getting Condition* materials through a series of focus groups with life science doctoral students (*n* = 26 across 6, 5-6 person groups). These students were selected from other universities (*n* = 16) across the United States to avoid contaminating the experimental sample for our RCT and to enhance the generalizability of the *Perspective-getting Condition* materials. We emailed the study information to students in their second to fourth year of life science doctoral programs. Students were again asked to provide detailed feedback on the *Perspective-getting Condition* materials. Specifically, participants were tasked with reading the materials and highlighting areas that elicited a positive or negative reaction, and to identify language that was off-putting or strange. Because of the large number of quotations in the *Perspective-getting Condition* across the two sessions, each focus group only reviewed a subset of the quotations in the materials. After each focus group, we iteratively revised the *Perspective-getting Condition* materials to remove language that was perceived as inauthentic or that invoked a strong negative reaction. We then tested the revised materials in the next focus group. Participants were offered a $25 gift card for participating in a focus group.

For the *Control Condition* materials, we sought feedback from six biology education researchers. These researchers were graduate students and postdoctoral researchers who had formal training in a combination of fields including biology, biology education, and psychology. We sought their feedback because collectively they were familiar with the norms and practices of mentoring relationships in STEM fields, had expertise in science education, and they were familiar with existing scholarship on mentorship. Their feedback helped us discover any aspects of the *Control Condition* materials that were perceived as inauthentic, distracting, or otherwise elicited a negative response. We revised the *Control Condition* content, quotations, and format based on their feedback.

**Intervention Adherence**

We verified intervention adherence (i.e., the extent to which students completed materials as intended) using several pieces of evidence. First, we assessed whether students completed the study activities at each timepoint. Of the 155 students who met the inclusion criteria, 149 students (96%) completed all Session 1 activities, and 134 students (86%) completed all Session 2 activities. Second, we assessed whether students engaged sufficiently with the materials by measuring the amount of time participants spent reading and completing the *Perspective-getting Condition* materials in their entirety, including the survey items. In estimating engagement time, we excluded 9 participants in Session 1 and 12 participants in Session 2 who had the materials open for >4 hours since these responses likely represent inactivity and not sustained engagement. On average, participants spent 29 minutes ($SD$ = 24 minutes) completing Session 1 and 38 minutes ($SD$ = 81 minutes) completing Session 2. Finally, we evaluated whether students recalled a negative interaction with their research mentor for which they might blame themselves, thus presenting a target for the intervention. Participant were asked to recall and write about a negative or unpleasant experience with their mentor in 2-3 sentences before they responded to the locus of causality scale. One researcher with expertise in negative mentoring experiences read each response to confirm reference to a negative interaction with their research mentor (Tuma et al., 2021). For the most part, participants described negative interactions with their research mentors (83% in Session 1 and 84% in Session 2) prior to responding about their locus of causality attributions. Thus, we concluded that students adhered to the materials largely as intended.

### Scales used to measure outcome variables

#### Research self-efficacy

Kardash, C. M. (2000). Evaluation of undergraduate research experience: Perceptions of undergraduate interns and their faculty mentors. *Journal of educational psychology*, *92*(1), 191.

**Instructions:** Please rate your confidence in your ability to perform the following tasks.
**Response Scale:** 1 = not at all confident; 2 = not very confident; 3 = somewhat confident; 4 = fairly confident; 5 = very confident; 6 = Prefer not to respond

| | |
|---|---|
| **Research self-efficacy** | 1. Understand contemporary concepts in your field. |
| | 2. Make use of the primary scientific research literature in your field (e.g., journal articles) |
| | 3. Identify a specific research question for investigation based on the research in your field |
| | 4. Formulate a research hypothesis based on a specific question |
| | 5. Design an experiment or theoretical test of the hypothesis |
| | 6. Understand the importance of "controls" in research |
| | 7. Observe and collect data |
| | 8. Statistically analyze data |
| | 9. Interpret data by relating results to the original hypothesis |
| | 10. Reformulate your original research hypothesis (as appropriate) |
| | 11. Relate the results to the "bigger picture" in your field |
| | 12. Orally communicate the results of research projects |
| | 13. Write a research paper for publication |
| | 14. Think independently |

**Locus of causality attributions for negative interactions**

McAuley, E., Duncan, T. E., & Russell, D. W. (1992). Measuring causal attributions: The revised causal dimension scale (CDSII). *Personality and Social Psychology Bulletin*, *18*(5), 566-573.

**Instructions:** Recall the last time you experienced any kind of tension or disagreement with your primary research advisor. This could be anything that happened with your advisor that made you feel frustrated, discouraged, or otherwise negatively in some way. Feel free to write about a new experience or the same one you wrote about last time.

In 2-3 sentences, explain the negative experience and how it made you feel.

*As a reminder, your responses will be kept completely confidential within the research team – no one outside the research team will see your responses.*

*(Text response)*

Think about the reason or reasons for the unpleasant experience you just wrote about. The items below concern your impressions or opinions of the cause or causes of your unpleasant experience. Select one scale-point for each of the following questions.

| Are the **cause(s)**: | | | | | | | | | | | |
|---|---|---|---|---|---|---|---|---|---|---|---|
| 1. | Something that reflects an aspect of the situation | 1 | 2 | 3 | 4 | 5 | 6 | 7 | 8 | 9 | Something that reflects an aspect of yourself |
| 2. | Outside of you | 1 | 2 | 3 | 4 | 5 | 6 | 7 | 8 | 9 | Inside of you |
| 3. | Something about others | 1 | 2 | 3 | 4 | 5 | 6 | 7 | 8 | 9 | Something about you |

**Mentoring relationship satisfaction**

**Item:** Overall, how satisfied are you with the quality of your relationship with your graduate mentor?
**Response Scale:** 1 = very dissatisfied; 2 = dissatisfied; 3 = neither satisfied nor dissatisfied; 4 = satisfied; 5 = very satisfied; 6 = prefer not to respond.

**Scholarly productivity**

**Instructions:** We would like to know more about what projects you work on in graduate school.
**Response scale:** Text entry
**Items:**

- How many first author publications do you have?
- How many co-authored publications (not counting first author publications) do you have?
- How many conference talks/poster presentations have you delivered?

**Results of measurement model fitting: confirmatory factor analyses**

Prior to performing our substantive analyses, we performed confirmatory factor analysis (CFA) using the R software (R Core Team, 2016) and the 'lavaan' package (Rosseel, 2012) for our self-efficacy and locus of causality attributions measures. We used the Robust Maximum Likelihood (MLR) estimator to account for non-normality in our data. We used several goodness-of-fit indices to examine how well our measurement models reproduced their variance-covariance matrices. Specifically, we examined both incremental (e.g., the Comparative Fix Index and the Tucker-Lewis Index) and parsimonious (e.g., the Root Mean Square of Error of Approximation and the Standardized Root Mean Square Residual) fit measures. Guided by Hu and Bentler (1999), we interpret CFI values $\geq$ 0.90, a RMSEA value < 0.06, and a SRMR < 0.08 as indicators of good model fit.

*Self-efficacy*
We ran a unidimensional CFA with self-efficacy indicated by 14 items, which indicated model misfit (Table S2). Across all four time points the CFI and TLI values were below the traditional cutoff of > 0.90 and the RMSEA was greater than 0.09. These results suggest that the unidimensional model have mis specified factor loadings for some of the indicators in the scale. To determine the potential underlying factor structure, we ran an EFA using principal axis factoring with an oblimin rotation on the self-efficacy scale from the Basline measure. Both the scree plot (Figure S1) and parallel analysis (Figure S2) indicated that a two-factor solution would best describe the data. The items from the two-factor solution all had loadings > 0.50, with the exception of two items which exhibited cross factor loadings (SE #7, *"Observe and collect data,"* and SE #10, "*Reformulate your original research hypothesis (as appropriate").*

We proceeded with splitting the self-efficacy scale into a two-factor model and dropped both items with cross loading from the measurement model. These modifications to the measurement model improved model fit (Table S3). We examined the content of the items in each factor, which appeared to reflect self-efficacy that was specific to two differences phases of research. The first factor related to self-efficacy that was specific to how to design and carry out investigations, which we termed "*investigation self-efficacy.*" The second factor related to self-efficacy regarding how to make sense of and communicate research results, which we termed "*interpretation self-efficacy.*" Thus, our substantive analyses deviated from our preregistration because our CFA results indicated our self-efficacy measure was best indicated as two separate outcome variables instead of one[1].

*Locus of causality attributions*
We ran a one-factor CFA model for our attributions scale indicated by 3 items. The model fit statistics indicated acceptable fit (Table S4). Therefore, we opted to proceed without adjusting the measurement model.

[1] The self-efficacy measure by Kardash (2000) was originally developed for use with undergraduate researchers and was adapted for use with doctoral students in the present study. Kardash (2000) reports evidence of reliability but modest evidence of the measure's validity. Furthermore, Kardash (2000) notes "The size of the sample in this study precluded a factor analysis of the instrument, although such an analysis should be undertaken in the future to provide further information regarding the instrument's psychometric properties and underlying constructs" (page 200).

**Table S2: Initial measurement model fit for self-efficacy**
*Note:* B = Baseline, P = Postintervention, L = Long-term evaluation

| Model | CFI | TLI | RMSEA | RMSEA 90% CI | SRMR | $\chi^2$ | df |
|---|---|---|---|---|---|---|---|
| Self-efficacy B | 0.80 | 0.77 | 0.13 | 0.11 – 0.15 | 0.07 | 251 | 77 |
| Self-efficacy P | 0.90 | 0.88 | 0.09 | 0.07 – 0.11 | 0.06 | 149 | 77 |
| Self-efficacy L | 0.83 | 0.80 | 0.12 | 0.10 – 0.14 | 0.08 | 188 | 77 |

### Figure S1: Scree plot for self-efficacy Baseline

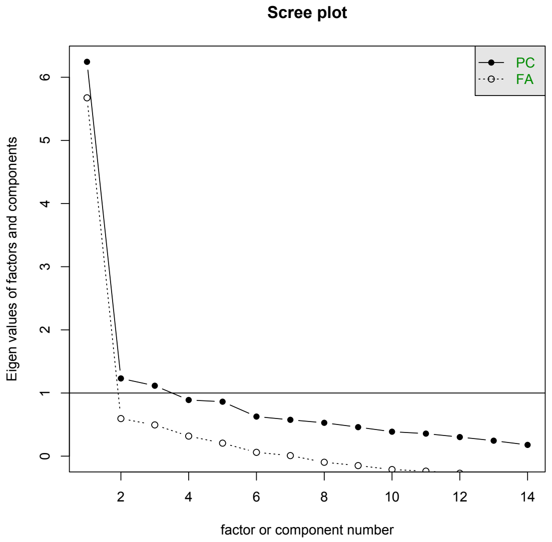


### Figure S2: Parallel analysis for self-efficacy Baseline

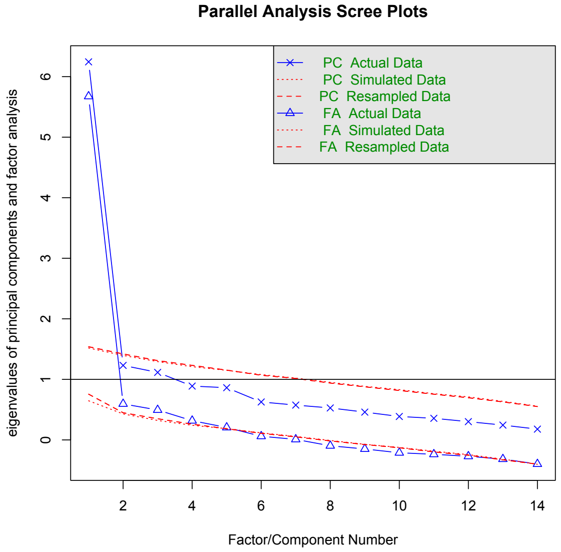

**Table S3: Measurement model fit for two factor model of self-efficacy**
*Factor 1:* Investigation self-efficacy indicated by items #1, 2, 3, 4, 5, and 6 (item 7 removed)
*Factor 2:* Interpretation self-efficacy indicated by items #8, 9, 11, 12, 13, and 14 (item 10 removed)
*Note:* B = Baseline, P = Postintervention, L = Long-term evaluation

| Model | CFI | TLI | RMSEA | RMSEA 90% CI | SRMR | χ2 | *df* |
|---|---|---|---|---|---|---|---|
| Self-efficacy B | 0.89 | 0.87 | 0.09 | 0.07 – 0.12 | 0.06 | 142.0 | 53 |
| Self-efficacy P | 0.93 | 0.91 | 0.08 | 0.06 – 0.11 | 0.06 | 109.4 | 53 |
| Self-efficacy L | 0.88 | 0.86 | 0.11 | 0.08 – 0.14 | 0.07 | 124.0 | 53 |

**Table S4: Measurement model fit for locus of causality attributions**

| Model | CFI | TLI | RMSEA | RMSEA 90% CI | SRMR | $\chi^2$ | *df* |
|---|---|---|---|---|---|---|---|
| Locus of causality attributions B | 1.00 | 1.00 | 0.00 | 0.00 – 0.00 | 0.00 | 72.8 | 3 |
| Locus of causality attributions B | 1.00 | 1.00 | 0.00 | 0.00 – 0.00 | 0.00 | 109.6 | 3 |
| Locus of causality attributions L | 1.00 | 1.00 | 0.00 | 0.00 – 0.00 | 0.00 | 115.2 | 3 |

**Concordance with Preregistered Analysis Plan**

We followed the analytical plan noted in the preregistration (https://osf.io/dfme7/?view_only=92678c7368a5446093bb263eeffe632d) with the following two minor deviations. First, because our self-efficacy measure did not exhibit sufficient fit to its proposed unidimensional factor structure as a 14-item measure (see Supplemental Materials for further details), we modeled self-efficacy as two separate variables (i.e., investigation self-efficacy and interpretation self-efficacy). Second, we decided before conducting analyses to analyze the data using regression instead of ANCOVA, and to use MPlus for analysis with full information maximum likelihood estimation for missing data instead of imputation, though we used the same model as we had proposed using originally.

**Examining Pre-Baseline Differences between the *Perspective-getting Condition* & Control Condition**

We first tested for significant differences in our five dependent variables and socio-demographic characteristics reported between the conditions at Baseline. To test for potential differences in continuous variables, we regressed initial levels of investigation self-efficacy, interpretation self-efficacy, locus of causality attributions, relationship satisfaction, and scholarly productivity on condition (intervention = +1, control = 0), controlling for year in program (1-4). For categorical variables, we conducted chi-square tests to examine whether the proportion of students in each categorical group (gender, race, first-generation status, and year in program) was approximately equal across conditions. The results revealed no significant pre-intervention differences between the *Perspective-getting Condition* and *Control Condition* for all variables.

**Correlations and Descriptive Statistics of Variables**

Doctoral students in our study reported high levels of investigation self-efficacy and more modest levels of interpretation self-efficacy. Their locus of causality attributions were generally more internal throughout the course of the study but exhibited large deviations. Students also reported high levels of satisfaction with their mentoring relationships on average. As expected, the two dimensions of research self-efficacy (i.e., investigation and interpretation) and scholarly productivity correlated significantly and positively with one another ($r$ = 0.26 – 0.41). Our measures of self-efficacy and locus of causality attributions had skewness that ranged from -0.77 to 0.61 and kurtosis that ranged from -0.88 and 1.08, thereby meeting assumptions for multivariate normality.
